# *Plasmodium vinckei* genomes provide insights into the pan-genome and evolution of rodent malaria parasites

**DOI:** 10.1101/2020.09.07.286369

**Authors:** Abhinay Ramaprasad, Severina Klaus, Olga Douvropoulou, Richard Culleton, Arnab Pain

## Abstract

**Background:** Rodent malaria parasites (RMPs) serve as tractable tools to study malaria parasite biology and host-parasite-vector interactions. *Plasmodium vinckei* is the most geographically widespread of the four RMP species collected in sub-Saharan Central Africa. Several *P. vinckei* isolates are available but relatively less characterized than other RMPs, thus hindering their use in experimental studies. We have generated a comprehensive resource for *P. vinckei* comprising of high-quality reference genomes, genotypes, gene expression profiles and growth phenotypes for ten *P. vinckei* isolates.

**Results:** The *P. vinckei* subspecies have diverged widely from their common ancestor and have undergone genomic structural variations. The subspecies from Katanga, *P. v. vinckei*, has a uniquely smaller genome, a reduced multigene family repertoire and is also amenable to genetic manipulation making it an ideal parasite for reverse genetics. Comparing *P. vinckei* genotypes reveals region-specific selection pressures particularly on genes involved in mosquito transmission. The erythrocyte membrane antigen 1 and *fam-c* families have expanded considerably among the lowland forest-dwelling *P. vinckei* parasites. Genetic crosses can be established in *P. vinckei* but are limited at present by low transmission success under the experimental conditions tested in this study.

**Conclusions:** *Plasmodium vinckei* isolates display a large degree of phenotypic and genotypic diversity and could serve as a resource to study parasite virulence and immunogenicity. Inclusion of *P. vinckei* genomes provide new insights into the evolution of RMPs and their multigene families. Amenability to genetic crossing and genetic manipulation make them also suitable for classical and functional genetics to study *Plasmodium* biology.

## Background

Rodent malaria parasites (RMPs) serve as tractable models for experimental genetics and as valuable tools to study malaria parasite biology and host-parasite-vector interactions [1–4]. Between 1948 and 1974, several rodent malaria parasites were isolated from wild thicket rats (shining thicket rat - *Grammomys poensis* or previously known as *Thamnomys rutilans*, and woodland thicket rat - *Grammomys surdaster*) and infected mosquitoes in sub-Saharan Africa and were adapted to laboratory-bred mice and mosquitoes. The isolates were classified into four species, namely, *Plasmodium berghei, Plasmodium yoelii, Plasmodium chabaudi and Plasmodium vinckei*. *Plasmodium berghei* and *P. yoelii* are sister species forming the classical *berghei* group, whereas *P. chabaudi* and *P. vinckei* form the classical *vinckei* group of RMPs [5–7]. *Plasmodium chabaudi* has been used for studying drug resistance, host immunity and immunopathology in malaria [8–12]. *Plasmodium yoelii* and *P. berghei* are extensively used as tractable models to study liver and mosquito stages of the parasite [13, 14]. Efficient transfection techniques [15–18] have been established in all three RMPs and they are widely used as *in vivo* model systems for large scale functional studies [19–22]. Reference genomes for these three RMP species are available [23, 24]. Recently, the quality of these genomes has been significantly improved using next-generation sequencing [11, 25, 26].

*P. vinckei* is the most geographically widespread RMP species, with isolates collected from many locations in sub-Saharan Africa (Figure 1A). Subspecies classifications were made for 18 *P. vinckei* isolates in total based on parasite characteristics and geographical origin, giving rise to five subspecies; *P. v. vinckei* (Democratic Republic of Congo), *P. v. petteri* (Central African Republic), *P. v. lentum* (Congo Brazzaville), *P. v. brucechwatti* (Nigeria) and *P. v. subsp.* (Cameroon) [27–32]. Blood, exo-erythrocytic and sporogonic stages of a limited number of isolates of the five subspecies have been characterized; *P. v. vinckei* line 67 or CY [33], *P. v. petteri* line CE [28], *P. v. lentum* line ZZ [29, 31], *P. v. brucechwatti* line 1/69 or DA [30, 34] and several parasite lines of *P. v. subsp.* [35]. Enzyme variation studies [5, 36] and multi-locus sequencing data [6, 7] have indicated that there is significant phenotypic and genotypic variation among *P. vinckei* isolates.

**Figure 1.**
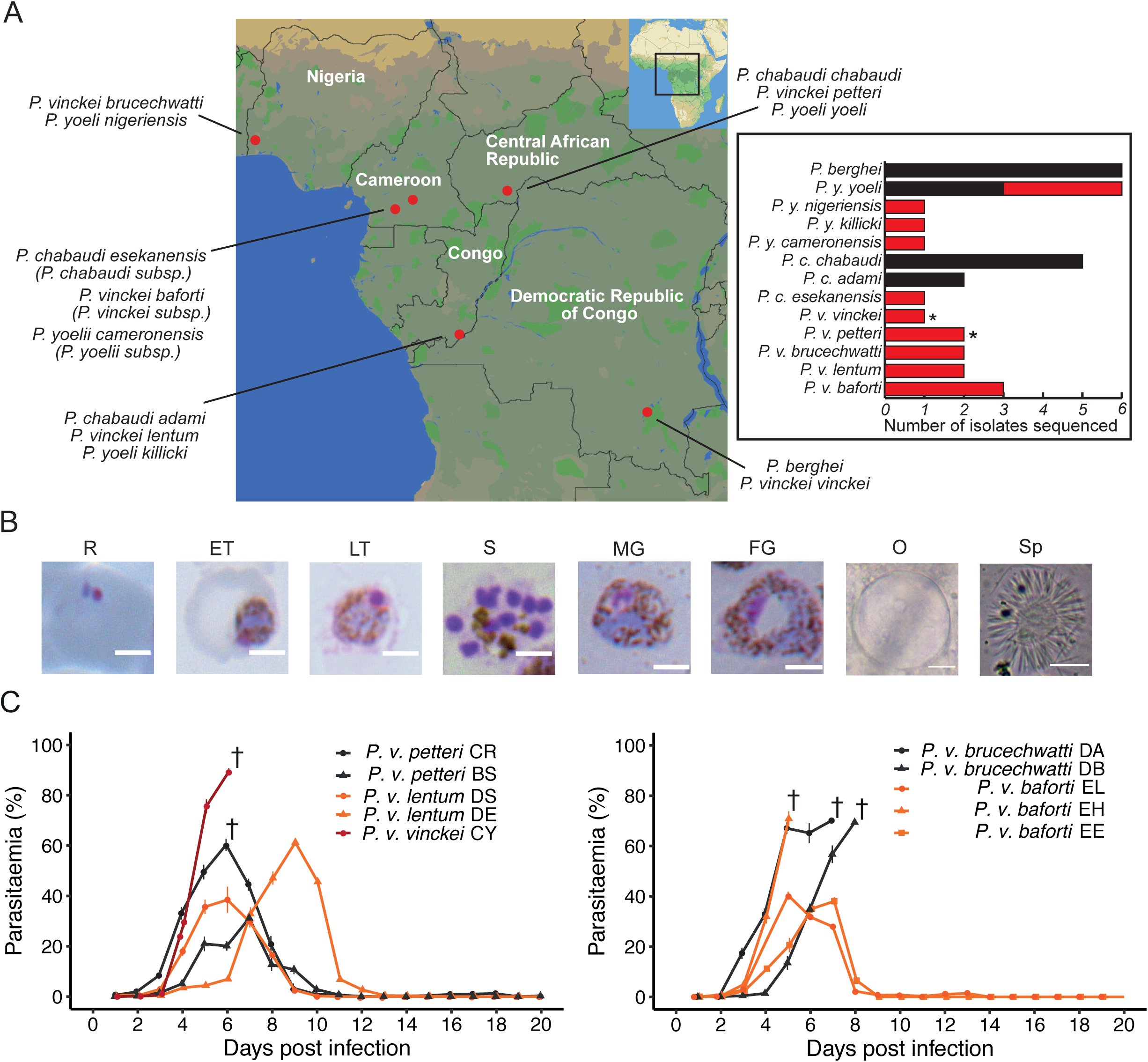
*Plasmodium vinckei* parasites and their phenotypic characteristics. A) Rodent malaria parasite species and subspecies and the geographical sites in sub-Saharan Africa where from which they were isolated (modified from [1]). *Plasmodium vinckei* is the only RMP species to have been isolated from five different locations. Inset: To date, several RMP isolates have been sequenced (black) to aid research with rodent malaria models. Additional RMP isolates have been sequenced in this study (red) to cover all subspecies of *P. vinckei* and further subspecies of *Plasmodium chabaudi* and *Plasmodium yoelii*. B) Morphology of different life stages of *P. vinckei baforti* EL. R: Ring, ET: early trophozoite, LT: Late trophozoite, S: Schizont, MG: Male gametocyte, FG: Female gametocyte, O: oocyst and Sp: Sporozoite. *Plasmodium vinckei* trophozoites and gametocytes are morphologically distinct from other RMPs due to their rich haemozoin content (brown pigment). C) Parasitaemia of ten *P. vinckei* isolates (split into two graphs for clarity) during infections in mice (n=5) for a 20-day duration. † denotes host mortality. *Plasmodium vinckei* isolates show significant diversity in their virulence phenotypes.

The rodent malaria parasites isolated from Cameroon in 1974 by J. M. Bafort are currently without subspecies names, being designated as *P. yoelii subsp.*, *P. vinckei subsp.* and *P. chabaudi subsp.*. We now present the full genome sequence data of isolates from these subspecies and show they form distinct clades within their parent species. Therefore, we propose the following subspecies names; *Plasmodium yoelii cameronensis*, from the country of origin; *Plasmodium vinckei baforti*, after J. M. Bafort, the original collector of this subspecies; and *Plasmodium chabaudi esekanensis*, from Eséka, Cameroon, the town from the outskirts of which it was originally collected.

Very few studies have employed *P. vinckei* compared to the other RMP species despite the public availability of several *P. vinckei* isolates (http://www.malariaresearch.eu/content/rodent-malaria-parasites). *Plasmodium v. vinckei* v52 and *P. v. petteri* CR have been used to study parasite recrudescence [37], chronobiology [38] and artemisinin resistance [39]. They are also the only isolates for which draft genome assemblies with annotation are available as part of the Broad Institute *Plasmodium* 100 Genomes initiative (https://www.ncbi.nlm.nih.gov/bioproject/163123).

A high-quality reference genome for *P. vinckei* and detailed phenotypic and genotypic data are lacking for the majority of *P. vinckei* isolates hindering wide-scale adoption of this RMP species in experimental malaria studies.

We now present a comprehensive genome resource for *P. vinckei* comprising of high-quality reference genomes for five *P. vinckei* isolates (one from each subspecies) and describe the genotypic diversity within the *P. vinckei* clade through the sequencing of five additional *P. vinckei* isolates (see Figure 1A inset). With the aid of high-quality annotated genome assemblies and gene expression data, we evaluate the evolutionary patterns of multigene families across all RMPs and within the subspecies of *P. vinckei*.

We also describe the growth and virulence phenotypes of these isolates and show that *P. vinckei* is amenable to genetic manipulation and can be used to generate experimental genetic crosses.

Furthermore, we sequenced the whole genomes of seven isolates of the subspecies of *P. chabaudi* (*P. c. esekanensis*) and *P. yoelii* (*P. y. yoelii*, *P. y. nigeriensis*, *P. y. killicki* and *P. y. cameronensis*) in order to resolve evolutionary relationships among RMP isolates.

The data presented here enable the use of the *P. vinckei* clade of parasites for laboratory-based experiments driven by high-throughput genomics technologies and will significantly expand the number of RMPs available as experimental models to understand the biology of malaria parasites.

## Results

### *Plasmodium vinckei* isolates display extensive diversity in virulence

We followed the infection profiles of ten *P. vinckei* isolates in CBA/J mice (five biological replicates per group) to study their virulence traits. Some of these isolates were available as uncloned lines and so were first cloned by limiting dilution (Additional File 1). As reported previously [40], *P. vinckei* parasites are morphologically indistinguishable from each other, prefer to invade mature erythrocytes, are largely synchronous during blood stage growth and display a characteristically rich abundance of haemozoin crystals in their trophozoites and gametocytes (Figure 1B).

Parasitaemia was determined daily to measure the growth rate of each isolate and host RBC density and weight were measured as indications of “virulence” (harm to the host) (Figure 1C, Additional file 2 and 3).

The *P. v. vinckei* isolate *Pvv*CY, was highly virulent and reached a parasitaemia of 89.4% ± 1.4 (standard error of mean; SEM) on day 6 post inoculation of 1 × 10^6^ blood stage parasites intravenously, causing host mortality on that day. Both strains of *P. v. brucechwatti*, *Pvb*DA and *Pvb*DB, were virulent and killed the host on day 7 or 8 post infection (peak parasitaemia of around 70%). The *P. v. lentum* parasites *Pvl*DS and *Pvl*DE, were not lethal and were eventually cleared by the host immune system, with *Pvl*DS’s clearance more prolonged than that of *Pvl*DE (parasitaemia clearance rates; *Pvl*DS = 10.35 %day^-1^; SE = 1.105; p-value of linear fit =0.0025; *Pvl*DE = 16.46 %day^-1^; SE = 3.873; p-value =0.023). The *P. v. petteri* isolates *Pvp*CR and *Pvp*BS reached peak parasitaemia along similar timelines (6-7 dpi), but *Pvp*CR was virulent (peak parasitaemia = 60.35 % ± 2.38 on day 6) and could sometimes kill the host while *Pvp*BS maintained a mild infection.

Of the three isolates of *P. vinckei baforti*, *Pvs*EL and *Pvs*EE were similar in their growth profiles and their perceived effect on the host, while in contrast, *Pvs*EH was highly virulent, causing host mortality at day 5, the earliest among all *P. vinckei* parasites.

RBC densities reduced during the course of infection proportionally to the rise in parasitaemia in all the *P. vinckei* infection profiles studied. There were differences, however, in the patterns of host weight loss. Mild infections by *P. v. lentum* isolates (maximum weight loss in *Pvl*DE = 0.43 mg ± 0.41 and *Pvl*DS = 1.77 mg ± 0.38), *P. v. petteri* BS (0.58 mg ± 0.22) and *P. v. baforti* EE (1.66 mg ± 0.31, 0.09 mg ± 0.56) did not cause any significant weight loss in mice, whereas the virulent strains, *P. v. petteri* CR (4.04 mg ± 0.18), *P. v. brucechwatti* isolates (*pvb*DA = 3.5 mg ± 0.39 and *pvb*DB = 2.05 mg ± 1.68) caused around a 20% decrease in weight. Virulent strains *Pvv*CY (1.74 mg ± 0.15) and *Pvs*EH (0.52 mg ± 0.13) did not cause any significant weight loss during their infection before host death occurred.

### *Plasmodium vinckei* reference genome assembly and annotation

High-quality reference genomes for five *P. vinckei* isolates, one from each subspecies; *P. v. vinckei* CY (*Pvv*CY), *P. v. brucechwatti* DA (*Pvb*DA), *P. v. lentum* DE (*Pvl*DE), *P. v. petteri* CR (*Pvp*CR) and *P. v. baforti* EL (*Pvs*EL) were assembled from single-molecule real-time (SMRT) sequencing. PacBio long reads of 10-20 kilobases (kb) and with a high median coverage of >155X across the genome (Additional File 1) enabled *de novo* assembly of each of the 14 chromosomes as single unitigs (high confidence contig) (see Table 1). PacBio assembly base call errors were corrected using high-quality 350bp and 550bp insert PCRfree Illumina reads. A small number of gaps remain in the assemblies, but these are mainly confined to the apicoplast genomes and to the *Pvs*EL and *Pvl*DE genomes that were assembled from 10kB-long PacBio reads instead of 20kB. The *Pvp*CR and *Pvv*CY assemblies, with each chromosome in one piece, are a significant improvement over their existing fragmented genome assemblies (available through PlasmoDB v.30).

**Table 1.** Genome assembly characteristics of five Plasmodium vinckei reference genomes. AT-rich *P. vinckei* genomes are 19.2 to 19.5 megabasepairs (Mbps) long except for *Pvv*CY which has a smaller genome size of 18.3 Mb, similar to *Plasmodium berghei.* PacBio long reads allowed for chromosomes to be assembled as gapless unitigs with a few exceptions. Number of genes include partial genes and pseudogenes. Copy numbers of the ten multigene families differ between the *P. vinckei* subspecies (*ema1*, erythrocyte membrane antigen 1, *etramp*, early transcribed membrane protein, *hdh*, haloacid dehalogenase-like hydrolase, *lpl*, lysophospholipases, *p235*, reticulocyte binding protein, *pir*, *Plasmodium* interspersed repeat protein).

| Nuclear genome features | *P. vinckei vinckei* CY | *P. vinckei brucechwatti* DA | *P. vinckei lentum* DE | *P. vinckei petteri* CR | *P. vinckei subsp. EL* |
| --- | --- | --- | --- | --- | --- |
| Genome Size (Mb) | 18.34 | 19.17 | 19.31 | 19.39 | 19.50 |
| G+C content (%) | 22.94 | 23.17 | 23.33 | 23.19 | 23.16 |
| Gaps within assembly | 0 | 0 | 7 | 0 | 11 |
| Genes | 5,042 | 5,238 | 5,249 | 5,310 | 5,307 |
| Pseudogenes | 19 | 40 | 29 | 36 | 36 |
| *ema1* | 7 | 20 | 19 | 15 | 21 |
| *etramp* | 12 | 12 | 11 | 13 | 12 |
| *fam-a* | 87 | 187 | 201 | 188 | 207 |
| *fam-b* | 28 | 38 | 41 | 40 | 44 |
| *fam-c* | 20 | 59 | 63 | 64 | 65 |
| *fam-d* | 6 | 13 | 27 | 21 | 21 |
| *hdh* | 7 | 9 | 12 | 13 | 14 |
| *lpl* | 25 | 23 | 24 | 23 | 17 |
| *p235* | 5 | 8 | 7 | 6 | 8 |
| pir | 178 | 209 | 210 | 272 | 247 |

*Plasmodium vinckei* genome sizes range from 19.2 to 19.5 Mb except for *Pvv*CY which has a smaller genome size of 18.3 Mb, similar to that of *P. berghei* (both isolates are from the same Katanga region). While we were not able to resolve the telomeric repeats at the ends of some of the chromosomes, all the resolved telomeric repeats had the RMP-specific sub-telomeric repeat sequences CCCTA(G)AA. The mitochondrial and apicoplast genomes were ∼6Kb and ∼30 kb long respectively, except for the apicoplast genomes of *Pvp*CR and *Pvs*EL for which we were able to resolve only partial assemblies due to low read coverage (see Additional File 4).

Gene models were predicted by combining multiple lines of evidence to improve the quality of those predictions. These include publicly available *P. chabaudi* gene models, *de novo* predicted gene models and transcript models from strand-specific RNA-seq data of different blood life cycle stages. Consensus gene models were then manually corrected through comparative genomics and visualization of mapped RNAseq reads. As a result, we annotated 5,073 to 5,319 protein-coding genes, 57-67 tRNA genes and 40-48 rRNA genes in each *P. vinckei* genome. Functional and orthology analyses with the predicted *P. vinckei* proteins showed that the core genome content in *P. vinckei* parasites is highly conserved among the species and are comparable to other rodent and primate malaria species.

### *Plasmodium vinckei* genome assemblies reveal novel structural variations

Comparative analysis of *P. vinckei* and other RMP genomes shows that *P. vinckei* genomes exhibit the same high level of synteny seen within RMP genomes, but with a number of chromosomal rearrangements. These events can be identified by breaks in synteny (synteny breakpoints-SBPs) observed upon aligning and comparing genome sequences.

We aligned *P. vinckei* and other RMP genomes to identify synteny blocks between their chromosomes. Similar to previous findings in RMP genomes [26, 41] (Additional file 12A), we observed large scale exchange of material between non-homologous chromosomes, namely three reciprocal translocation events and one inversion (Figure 2A, Additional File 5 and 12). A pan-*vinckei* reciprocal translocation of ∼0.6Mb (with 134 genes) and ∼0.4 Mb (with 99 genes) long regions between chromosomes VIII and X was observed between *P. vinckei* and *P. berghei* (whose genome closely resembles that of the putative RMP ancestor [41]). Within the *P. vinckei* subspecies, two reciprocal translocations separate *P. v. petteri* and *P. v. baforti* from the other three subspecies. One pair of exchanges (∼1 Mb and ∼0.55 Mb) was observed between chromosomes V and XIII, and another smaller pair (∼150Kb and ∼70Kb) between chromosomes V and VI. These events have left the Chromosomes V of *Pvv*CY-*Pvb*DA-*Pvl*DE and *Pvp*CR-*Pvs*EL groups with only a ∼0.15 Mb region of synteny between them, consisting of 48 genes while the remaining 304 genes have been rearranged with chromosome VI and XII.

**Figure 2.**
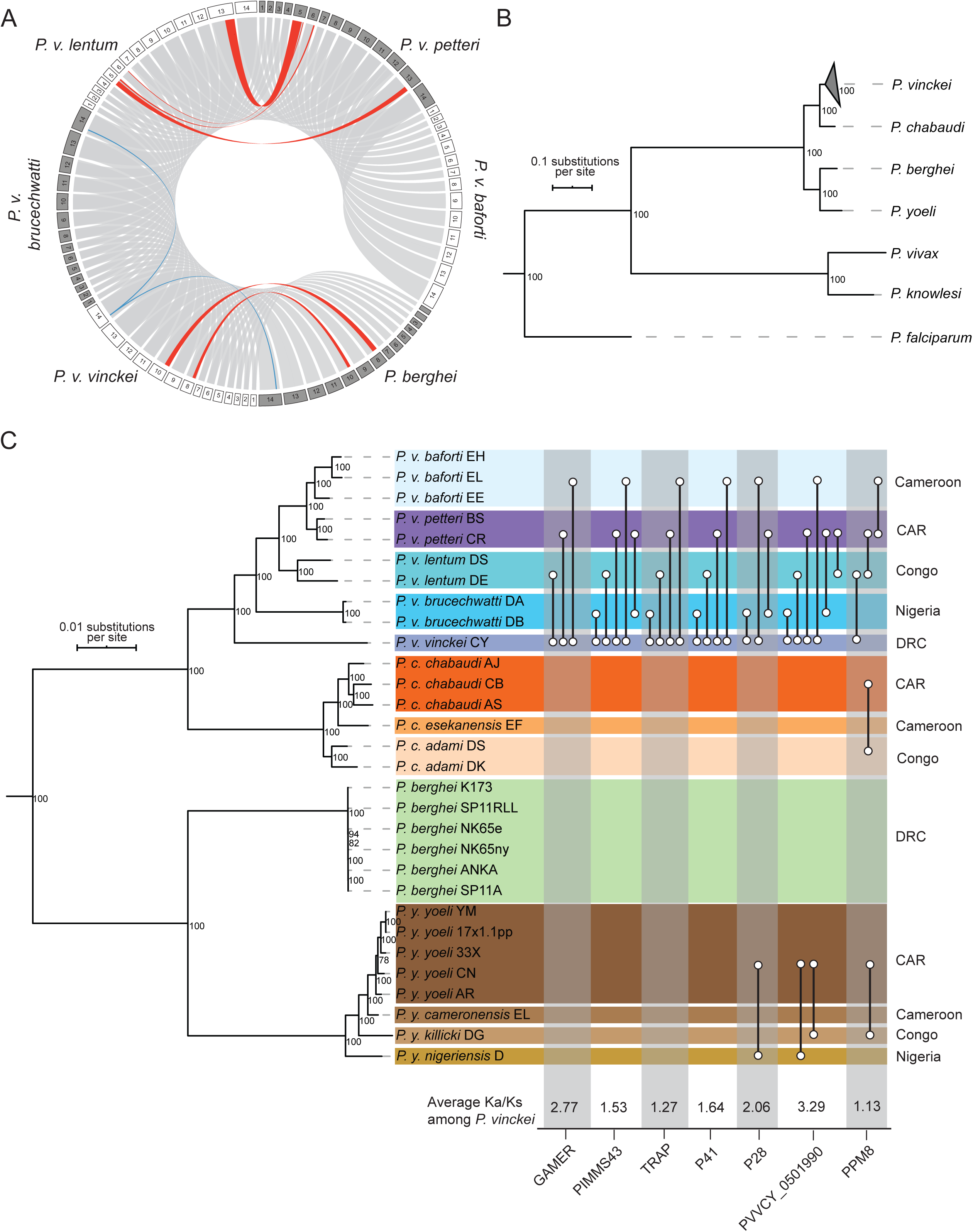
Structural variations and genotypic diversity among *Plasmodium vinckei* parasites. A) Chromosomal rearrangements in *P. vinckei* parasites. Pairwise synteny was assessed between the five *P. vinckei* subspecies and *Plasmodium berghei* (to represent the earliest common RMP ancestor). The 14 chromosomes of different RMP genomes are arranged as a Circos plot and the ribbons (grey) between them denote regions of synteny. Three reciprocal translocation events (red) and one inversion (blue) accompany the separation of the different *P. vinckei* subspecies. A pan-*vinckei* reciprocal translocation between chromosomes VIII and X was observed between *P. vinckei* and other RMP genomes. Within the *P. vinckei* subspecies, two reciprocal translocations, between chromosomes V and XIII, and between chromosomes V and VI, separate *Plasmodium vinckei petteri* and *P. v. baforti* from the other three subspecies. A small inversion of ∼100 kb region in chromosome 14 has occurred in *Pvv*CY alone. B) Maximum likelihood phylogeny of different RMP species with high-quality reference genomes based on protein alignment of 3,920 one-to-one orthologs (bootstrap values of each node are shown). Genomes of three human malaria species- *Plasmodium falciparum*, *Plasmodium vivax* and *Plasmodium knowlesi* were included in the analysis as outgroups. C) Maximum likelihood phylogenetic tree of all sequenced RMP isolates based on 1,010,956 high-quality SNPs (bootstrap values of each node are shown). There exists significant genotypic diversity among the *P. vinckei* isolates compared to the other RMPs. All *P. vinckei* subspecies have begun to diverge from their common ancestor well before sub-speciation events within *Plasmodium yoelii* and *Plasmodium chabaudi*. Genetic diversity within *P. v. petteri* and *P. v. baforti* isolates are similar to those observed within *P. yoelii* and *P. chabaudi* isolates while *P. v. lentum* and *P. v. brucechwatti* isolates have exceptionally high and low divergences respectively. Genes with significantly high Ka/Ks ratios in different subspecies-wise comparisons (as indicated by connector lines), the gene’s Ka/Ks ratio averaged across all indicated *P. vinckei* comparisons and geographical origin of the isolates are shown.

There also exists a small, *Pvv*CY-specific inversion of a ∼100 kb region in chromosome XIV. All the synteny breakage points (SBPs) were verified manually and were supported by PacBio read coverage ruling out the possibility of a misassembly at the breakpoint junctions. The SBPs in chromosomes V and VI were near rRNA units, loci previously described as hotspots for such rearrangement events [42, 43].

### A pan-RMP phylogeny reveals high genotypic diversity within the *P. vinckei* clade

In order to re-evaluate the evolutionary relationships among RMPs, we first inferred a well-resolved species-level phylogeny that takes advantage of the manually curated gene models in eight available high-quality RMP genomes representing all RMPs. A maximum-likelihood phylogeny tree was inferred through partitioned analysis using RAxML, of a concatenated protein alignment (2,281,420 amino acids long) from 3,920 single-copy, conserved core genes in eleven taxa (eight RMPs, *P. falciparum*, *P. knowlesi* and *P. vivax*; see Figure 2B and Additional File 6).

In order to assess the genetic diversity within RMP isolates, we sequenced additional isolates for four *P. vinckei* subspecies (*Pvb*DB, *Pvl*DS, *Pvp*BS, *Pvs*EH and *Pvs*EE), *P. yoelii yoelii* (*Pyy*33X, *Pyy*CN and *Pyy*AR), *P. yoelii nigeriensis* (*Pyn*D), *P. yoelii killicki* (*Pyk*DG), *P. yoelii subsp.* (*Pys*EL) and *P. chabaudi subsp.* (*Pcs*EF) (Additional File 1). This, along with existing sequencing data for 13 RMP isolates (from [26, 44]), were used to infer an isolate-level, pan-RMP maximum likelihood phylogeny based on 1,010,956 high-quality SNPs in non-subtelomeric genes that were called by mapping all reads onto the *Pvv*CY reference genome (Figure 2C and Additional File 7). Both phylogenies were well-resolved with robust 100% bootstrap support obtained for the amino-acid based phylogeny and 78% or higher bootstrap support for the SNPs-based phylogeny (majority-rule consensus tree criterion was satisfied at 50 bootstraps for both the phylogenies).

Both protein alignment-based and SNP-based phylogenies show significant divergence among the *P. vinckei* subspecies compared to the other RMPs. All *P. vinckei* subspecies have begun to diverge from their common ancestor well before sub-speciation events within *P. yoelii* and *P. chabaudi*.

A total of 521,934 polymorphic positions were found within the *P. vinckei* core coding regions consisting of 4,644 non-subtelomeric genes across *P. vinckei* isolates. The Katangan isolate, *P. v. vinckei*, has undergone significant divergence from the common *vinckei* ancestor and is the most diverged of any RMP subspecies sequenced to date. Number of SNPs ranged from 292,240 to 318,344 in pair-wise comparisons of isolates with *Pvv*CY, with 4,237 to 4,263 genes (out of the 4,644 core genes) having at least 5 SNPs. *Plasmodium v. brucechwatti* has also diverged significantly, while the divergence of *P. v. lentum* is comparable to that of *P. y. nigeriensis*, *P. y. kilicki* and *P. c. subsp.* from their respective putative ancestors. Genetic diversity within *P. v. petteri* and *P. v. baforti* isolates are similar to that observed within *P. yoelii* and *P. chabaudi* isolates while *P. v. lentum* and *P. v. brucechwatti* isolates have exceptionally high and low divergences respectively. Number of SNPs in pair-wise comparisons between the closest subspecies, *P. v. petteri* and *P. v. baforti*, ranged from 53,554 (*P. v. baforti* EE) to 69,454 (*P. v. baforti* EL) with 2,358 genes having at least 5 SNPs.

Our robust phylogeny based on a comprehensive set of genome-wide sequence variations confirms previous estimates of RMP evolution based on isoenzyme variation [5] and gene sequences of multiple housekeeping loci [6, 7], except for the placement of *P. y. nigeriensis* D which we show to be diverged earlier than *P. y. kilicki* DG (supported by a bootstrap value of 100).

### Molecular evolution within *P. vinckei* isolates

Using SNP data (Additional file 7), we then assessed the differences in selection pressure on the geographically diverse *P. vinckei* isolates by calculating the gene-wise Ka/Ks ratio as a measure of enrichment of non-synonymous mutations in a gene (signifying positive selection). We first compared the Katangan isolate (*Pvv*CY) from the highland forests in the DRC with the non-Katangan isolates from the lowland forests elsewhere.

We made pairwise comparisons of the four non-Katangan *P. vinckei* subspecies with *P. v. vinckei* which revealed several genes under significant positive selection (Figure 3C). Notably, we identified three genes involved in mosquito transmission, namely, a gamete-release protein (GAMER), a secreted ookinete protein (PIMMS43, previously known as PSOP25) and a thrombospondin-related anonymous protein (TRAP), featuring in all Katangan/non-Katangan subspecies comparisons.

**Figure 3.**
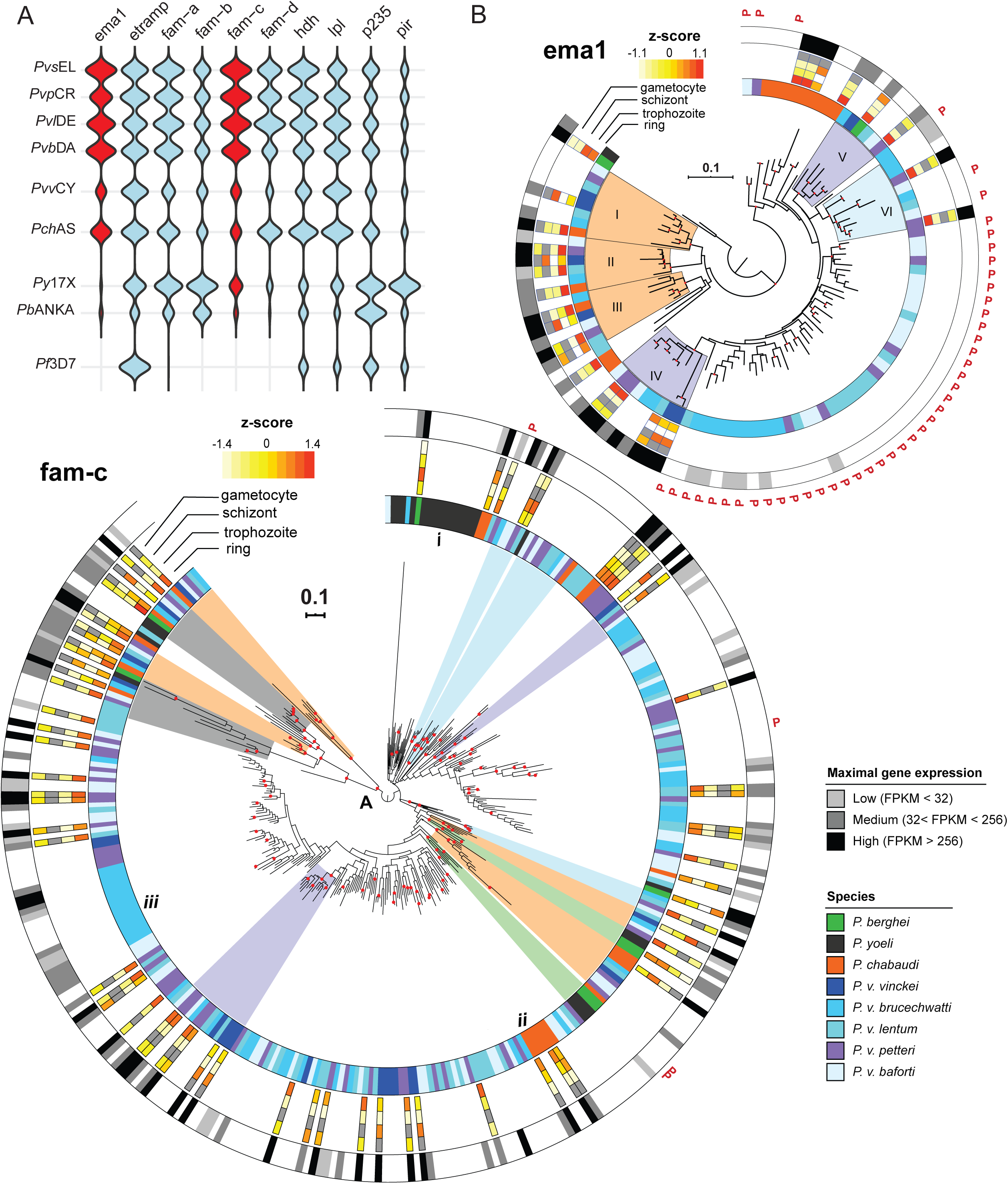
Sub-telomeric multigene family expansions in *Plasmodium vinckei* parasites. **A)** Violin plots show sub-telomeric multigene family size variations among RMPs and *Plasmodium falciparum*. The erythrocyte membrane antigen 1 and *fam-c* multigene families are expanded in the non-Katangan *P. vinckei* parasites (red). Apart from these families, multigene families have expanded in *P. vinckei* similar to that in *Plasmodium chabaudi*. The Katangan isolate *Pvv*CY (purple) has a smaller number of family members compared to non-Katangan isolates (orange) except for lysophospholipases, *p235* and *pir* gene families. B) Maximum Likelihood phylogeny of 99 *ema1* (top) and 328 *fam-c* (bottom) genes in RMPs. Branch nodes with good bootstrap support (> 70) are marked in red. The first coloured band denotes the RMP species to which the particular gene taxon belongs to. The heatmap denotes the relative gene expression among rings, trophozoite, schizont and gametocyte stages in the RMPs for which data are publicly available. Orange denotes high relative gene expression and white denotes low relative gene expression, while grey denotes lack of information. Gene expression was classified into three categories based on FPKM level distribution-High (black) denotes the top 25% of ranked FPKM of all expressed genes (FPKM > 256), Low (light grey) is the lower 25% of all expressed genes (FPKM < 32) and Medium level expression (grey;32<FPKM<256). “P” denotes pseudogenes. Four *vinckei*-group (*P. chabaudi* and *P. vinckei* subspecies) specific clades (Clades I-IV; orange), two *vinckei*-specific clade (Clade IV and V; purple) and one non-Katangan-specific clade (Clade VI; blue) can be identified within *ema1* family with strong gene expression, maximal during ring stages. Rest of the family’s expansion within non-Katangan *P. vinckei* isolates are mainly pseudogenes with weak transcriptional evidence. The fam-c gene phylogeny shows the presence of four distinctly distal clades (A) with robust basal support (96-100). Of the four clades, two are pan-RMP (grey) and two are *vinckei* group-specific (orange), each consisting of fam-c genes positionally conserved across the member subspecies. Most members of these clades show medium to high gene expression during asexual blood stages. Other well-supported clades can be classified as either *berghei* group-specific (two; green), *vinckei* group-specific (two; orange), *P. vinckei* - specific (two; purple) or non-Katangan *P. vinckei* -specific clades (three; blue). There is evidence of significant species-specific expansion with striking examples in *Plasmodium yoelii* (i), *P. chabaudi* (ii) and in *P. v. brucechwatti* (iii).

GAMER (PVVCY_1202630 being the representative ortholog in *P. v. vinckei*; PBANKA_1225400 in *P. berghei*) had high Ka/Ks values in all comparisons (except for *P. v. vinckei*-*P. v. brucechwatti*) and is essential for gamete egress [45]. PIMMS43 (PVVCY_1102000; PBANKA_1119200) and TRAP (PVVCY_1305250; PBANKA_1349800) showed high Ka/Ks values in all comparisons and are essential for ookinete evasion from mosquito immune system [46] and sporozoite infectivity of mosquito salivary glands and host hepatocytes [47] respectively.

Several exported proteins and surface antigens were also identified to have undergone positive selection. PVVCY_0100120 (PCHAS_0100651 being the gene ortholog in *P. chabaudi*) has a circumsporozoite-related antigen PFAM domain (PF06589) and is a conserved protein found in all RMPs except *P. berghei*. PVVCY_1200100 (PBANKA_1002600) is a merozoite surface antigen, p41 [48] that is secreted following invasion [49].

To assess presence of geographic location-specific selection pressures among the lowland forest isolates, *P. v. brucechwatti*, *P. v. lentum* and *P. v. baforti* were compared with *P. v. petteri* CR from the CAR. To see if similar selection pressures have acted on other RMP species, we also analysed the *P. yoelii* and *P. chabaudi* isolates from these regions that we had sequenced in this study.

Several exported and rhoptry-associated proteins were identified as been under positive selection in each comparison but in contrast to comparisons with *P. v. vinckei*, there was no overlap of positively selected genes among the non-Katangan isolates. However, we identified a conserved rodent malaria protein of unknown function (PVVCY_0501990; PBANKA_051950) that seems to be under significant positive selection with high Ka/Ks values (ranging from 2.14 to 4.39) in all *P. vinckei* comparisons except *P. v. petteri - P. v. baforti*. The *P. yoelii* ortholog of this protein was also positively selected among *P. y. yoelii, P. y. nigeriensis* and *P. y. killicki* but was not under selection within the *P. y. yoelii* isolates, signifying region-specific selection pressures.

A 28kDa ookinete surface protein (P28; PVVCY_0501540; PBANKA_0514900) seem to be under positive selection in the Nigerian *P. v. brucechwatti* as it features in both *P. v. vinckei - P. v. brucechwatti* and *P. v. brucechwatti - P. v. petteri* comparisons. The protein is also seen positively selected among corresponding Nigerian and Central African Republic *P. yoelii* isolates (*P. y. nigeriensis – P. y. yoelii*). A protein phosphatase (PPM8; PVVCY_0903370; PBANKA_0913400) has also undergone positive selection in all three RMP species between CAR and Congo isolates (*P. c. chabaudi- P. c. adami* comparison from [26]).

### Evolutionary patterns within the RMP multigene families

We were able to accurately annotate members of the ten RMP multigene families in the *P. vinckei* genomes owing to the well-resolved sub-telomeric regions in the Pacbio assemblies and manually curated gene models (see Table 1 and Additional file 8). *P. v. vinckei*, similar to its sympatric species, *P. berghei*, had a lower multigene family repertoire with copy numbers strikingly less than other *P. vinckei* subspecies (exceptions were the *pir*, *etramp* and lysophospholipase families). Multigene family sizes in the four non-Katangan *P. vinckei* subspecies were similar to *P. chabaudi* except for expansion in the *ema1* and *fam-c* multigene families (Figure 3A).

Next, we inferred maximum likelihood-based phylogenies for the ten multigene families, in order to identify structural differences amongst their members and to determine family evolutionary patterns across RMP species and *P. vinckei* subspecies (Figure 3B, Additional file 3 and 4). Overall, we identified robust clades (with bootstrap value >70) that fell into the following categories, i) pan-RMP, with orthologous genes from the four RMP species (dark grey), ii) *berghei* group, with genes from *P. berghei* and *P. yoelii* alone, iii) *vinckei* group, with genes from *P. chabaudi* and any or all *P. vinckei* subspecies, iv) *P. vinckei*, with genes only from *P. vinckei* subspecies and v) non-Katangan, with genes from all *P. vinckei* subspecies except *P. v. vinckei*.

In general, a high level of orthology was observed between *P. chabaudi* and *P. vinckei* genes forming several *vinckei* group clades (marked in orange in Figure 3) in contrast to more species-specific clades of paralogous genes being formed in *P. berghei* and *P. yoelii*. Thus, family expansions in *P. chabaudi* and *P. vinckei* seem to have occurred in the common *vinckei* group ancestor prior to speciation. We rebuilt the phylogenetic trees for *pir*, *fam-a* and *fam-b* families in order to see if the previously defined clades [25, 26] were also maintained in *P. vinckei*. Overall, we were able to reproduce the tree structures for the three families using the ML method with *P. vinckei* gene family members now added to them.

For *pirs*, we obtained four long-form and eight short-form clades as in [26] (Additional file 9, tree 10) albeit with lower bootstrap support, possibly due to our overly stringent automated trimming of the sequence alignment (see Methods). With a few exceptions, *P. vinckei pir* genes majorly populated two clades - L1 and S7 and a subclade S1g. These clades, previously shown to be *P. chabaudi*-dominant, hold equal or near equal proportions of *P. vinckei* pirs too. The only other *P. chabaudi*- dominant clade, L4, remains as a completely *P. chabaudi*-specific gene expansion. No *P. vinckei* species- or subspecies-specific clades are evident except for two subclades that could be inferred as *Pvv*CY-specific expansions within L1 and S7 (marked i and ii in Additional file 9, tree 10). Speciation of *P. vinckei* subspecies from their common ancestor seems to have been accompanied by gene gain in L1 and S7 clades and gene loss in S1g subclade. There is an almost linear increase of around 20 genes in *Pvv*CY, *Pvb*DA and *Pvl*DE in clade L1 pirs and a near doubling of clade S7 pirs in *Pvp*CR.

The *fam-a* and *fam-b* phylogenies (Additional file 9, tree 3 and 4 respectively) show that previously identified ancestral lineages [25] are maintained in *P. vinckei* too. The addition of *P. vinckei* genes resolved the ancestral clade of internal *fam-a* genes in chromosome 13 further into several well-supported *vinckei* group clades and a *berghei* group clade (marked as A in Additional file 9, tree 3). The 19 other *fam-a* clades and five *fam-b* clades consisting of positionally conserved orthologous genes are also conserved in *P. vinckei* (pan-RMP clades marked with * in Additional file 9, tree 3 and 4). The *fam-a* family has expanded in the non-Katangan *P. vinckei* subspecies through independent events of gene duplication in their common ancestor giving rise to several non-Katangan clades (marked as B in Additional file 9, tree 3). There is only a moderate *P. vinckei*-specific expansion in *fam-b* giving rise to three clades (marked as A in Additional file 9, tree 4) that includes *Pvv*CY genes too, pointing to gene duplications in the *P. vinckei* common ancestor. In both the phylogenies, species and subspecies-specific gene duplication events within the *vinckei* group are rare but do occur (marked as i-iv in Additional file 9, tree 3 and 4).

The *fam-d* multigene family is present as a single ancestral copy in *P. berghei* internally on chromosome IX but is expanded into a gene cluster in the same loci in *P. yoelii* (5 genes) and *P. chabaudi* (21 genes). Similar expansions have occurred in *P. vinckei* subspecies and phylogenetic analysis shows the presence of six robust clades within this family (Additional file 9, tree 6). Clade I is clearly the ancestral clade from which all other *fam-d* genes have been derived as it consists of the single *P. berghei* gene and its orthologs in other RMPs, positionally conserved to be the outermost gene of the *fam-d* cluster in each RMP.

While the *fam-d* family in *P. yoelii* is completely a product of paralogous expansion within Clade I, the *fam-d* families in the *vinckei* group seem to have expanded *via* five ancestral lineages forming clades II-VI. A subset of orthologs in Clade II (marked with * in Additional file 9, tree 6) are positionally conserved among *vinckei* group parasites, located immediately after the *fam-d* ancestral copy and could therefore represent the Clade II ancestral gene in the *vinckei* group common ancestor. *Pvv*CY has a smaller *fam-d* repertoire of 6 genes derived from only three of the five *vinckei* group lineages (Clade II, IV and VI), apart from the conserved ancestral copy.

ML-based trees for haloacid dehalogenase-like hydrolase (*hdh*), putative reticulocyte binding proteins (*p235*) and lysophospholipases (*lpl*) have generally well-resolved topologies with robust bootstrap support for their nodes and some clades contain syntenic orthologous genes (clades marked with * in Additional file 9, trees 7, 8 and 9) to member genes in *P. falciparum,* for example, *Pf*HAD2, *Pf*HAD3, *Pf*HAD4 and *Pf*RH6. Poor bootstrap support was obtained for the *etramp* tree (Additional file 9, tree 2), however clades were identified for some members including *uis3*, *uis4* and *etramp10.2*.

In order to assess the general level of expression of multi-gene family members in blood stage parasites, we superimposed blood-stage RNAseq data onto the phylogenetic trees. Life stage specific expression of multigene family members in five RMPs – *P. berghei* (rings, trophozoites, schizonts and gametocytes) [26], *P. chabaudi* and *P. v. vinckei* (rings, trophozoites and schizonts) [50], *P. v. petteri* and *P. v. lentum* (rings, trophozoites and gametocytes) and mixed blood stage expression levels in *P. yoelii* [44], *P. v. brucechwatti* and *P. v. baforti* were assessed (Additional File 10).

The level of gene transcription was designated low for genes with normalized FPKM (Fragments Per Kilobase of transcript per Million mapped reads) less than 36, medium if between 36 and 256 and high for genes with FPKM above 256. Both the levels and life stage specificity of gene expression within the various clades were generally conserved across the RMPs signifying that orthologs in structurally distinct clades might have conserved functions across the different RMPs. In general, the proportion of transcribed genes in all multigene families in *P. vinckei* was similar to that observed in *P. chabaudi* and slightly higher than *P. berghei* and *P. yoelii* (excluding the families with only one or two *P. berghei* or *P. yoelii* members).

### The erythrocyte membrane antigen 1 and *fam-c* sub-telomeric multigene families are expanded in non-Katangan *P. vinckei* parasites

The erythrocyte membrane antigen 1 (EMA1) was first identified and described in *P. chabaudi* and is associated with the host RBC membrane [51]. These genes encode for a ∼800 aa long protein and consist of two exons; a first short exon carrying a signal peptide followed by a longer exon carrying a PcEMA1 protein family domain (Pfam ID-PF07418). The gene encoding EMA1 is present only as a single copy in *P. yoelii* or as two copies in *P. berghei* but has expanded to 14 genes in *P. chabaudi*. We see similar gene expansions of 15 to 21 members in the four non-Katangan *P. vinckei* parasites (*Pvb*DA, *Pvl*DE, *Pvp*CR and *Pvs*EL; Figure 3A and Additional file 3). However, almost half of these genes are pseudogenes with a conserved SNP (C>A) at base position 14 that introduces a TAA stop codon (S5X) within the signal peptide region, followed by a few more stop codons in the rest of the gene (see Additional File 12B). Apart from one or two cases, the S5X mutation is found in all pseudogenes belonging to the *ema1* family and is *vinckei*-specific (it is not present in the single *P. chabaudi* pseudogene).

A ML-based phylogeny inferred for the 99 *ema1* genes was in general well-resolved with robust branch support for most nodes (see Figure 3B). Four distinct *vinckei* group-specific clades (Clade I to IV), two *vinckei*-specific clades (Clade IV and V) and a non-Katangan *P. vinckei* - specific clade (VI) with good basal support (bootstrap value of 75-100) were identified. Clade I-IV each consist of *ema1* genes positionally conserved across *P. chabaudi* and all five *P. vinckei* subspecies (in chromosomes I, VII, IX and X respectively) and are actively transcribed during blood stages.

Of the two *P. berghei ema1* genes, one forms a distal clade with the single *P. yoelii* gene while the other is paraphyletic within Clade IV, pointing to presence of two *ema1* loci in the common RMP ancestor, one of which was possibly lost during speciation of *P. yoelii*. All seven *Pvv*CY *ema1* genes are found within clades I to V with gene duplication events in Clade III and V.

Family expansion in *P. chabaudi* is mainly driven by gene duplication giving rise to *P. chabaudi*-specific clades. In contrast, family expansion within non-Katangan *P. vinckei* parasites is mainly driven by expansion of pseudogenized *ema1* genes (41 genes). Except for some *P. v. brucechwatti*-specific gene expansions, the pseudogenes do not form subspecies-specific clades suggesting that the expansion must have occurred in their non-Katangan *P. vinckei* common ancestor.

Gene expression data shows that the members of the *P. vinckei*-specific Clade IV are heavily transcribed during blood stages but most *ema1* pseudogenes that share ancestral lineage with Clade IV are very weakly transcribed. Taken together, a core repertoire of conserved *ema1* genes arising from 4-5 independent ancestral lineages are actively transcribed during blood stages of *P. vinckei* and *P. chabaudi*. The *ema1* multigene family expansion in *P. vinckei* is largely due to duplications of *ema1* pseudogenes, all carrying a S5X mutation and lacking transcription.

The *fam-c* proteins are exported proteins characterized by *pyst-c1* and *pyst-c2* domains, first identified in *P. yoelii* [42]. There is a considerable expansion of this family in the non-Katangan *P. vinckei* strains resulting in 59 to 65 members, twice that of *Pvv*CY and other RMPs. The *fam-c* genes are exclusively found in the sub telomeric regions and are composed of two exons and an intron, of which the first exon is uniformly 80 bps long (with a few exceptions). *fam-c* proteins are approximately 100-200 amino acids long, and more than one third of the proteins in *P. vinckei* contain a transmembrane domain (75.5%) and a signal peptide (88.9%) but most of them lack a PEXEL-motif (motif was detected in only 4% of the genes compared to 24% in other RMPs).

An ML-based tree of all *fam-c* genes in the eight RMP species shows the presence of four distinctly distal clades (marked as A in Figure 3B) with robust basal support (96-100). Two of them are pan-RMP and two are *vinckei* group-specific, each consisting of *fam-c* genes positionally conserved across the member subspecies (taking into account the genome rearrangement between chromosome V and VI within the *vinckei* clade). Most members of these clades show medium to high gene expression during asexual blood stages.

The remainder of the tree’s topology does not have good branch support (<70) with the exception of some terminal nodes, but it does demonstrate the significant expansion of this gene family within non-Katangan *P. vinckei* parasites (clades shaded in blue).

There is evidence of significant species- and subspecies-specific expansions with striking examples in *P. yoelii*, *P. chabaudi* and in *P. v. brucechwatti* (marked i, ii and iii in Figure 3B respectively), though they do not form well-supported clades. Most *fam-c* genes in *P. yoelii* seem to have originated from such independent *P. yoelii* - specific expansion events. *P. chabaudi* and *P. v. vinckei fam-c* genes are found more widely dispersed throughout the tree suggesting divergence of this family in the *vinckei* group common ancestor. On the other hand, subspecies-wise distinctions among the non-Katangan *P. vinckei fam-c* genes are less resolved as they form both paralogous and orthologous groups between the four subspecies with several ortholog pairs strongly supported by bootstrap values.

Thus, *fam-c* gene family expansion in the non-Katangan *P. vinckei* subspecies seems to have been driven by both gene duplications in their common ancestor and subspecies-specific gene family expansions subsequent to subspeciation. Around half of the *fam-c* genes have detectable transcripts in asexual or sexual blood stages. Most of the transcribed genes have medium (36<FPKM<256) or high-level expression (FPKM > 256) and blood-stage specific expression data for *P. chabaudi* and *P. v. vinckei* show peak transcription among the asexual blood stages at ring and schizont stages.

### Genetic crossing can be performed between *P. vinckei* isolates

The availability of several isolates within each *P. vinckei* subspecies with varying growth rates and wide genetic diversity makes them well-suited for genetic studies. Therefore, we attempted genetic crossing of the two *P. vinckei baforti* isolates, *Pvs*EH and *Pvs*EL, that displayed differences in their growth rates. Optimal transmission temperature and vector stages were initially characterized for *P. v. baforti* EE, EH and EL. Each isolate was inoculated into three CBA mice and on day 3 post infection, around 100 female *A. stephensi* mosquitoes were allowed to engorge on each mouse at different temperatures - 21°C, 23°C and 26°C. All three *P. v. baforti* isolates were able establish infections in mosquitoes at 23°C and 26°C, producing at least 50 mature oocysts on day 15 post-feed, but failed to transmit at 21°C (Figure 1A (a) and Additional file 6). Four to five oocysts of 12.5-17.5 µm diameter were observed at day 8 post-feed in the mosquito midgut and around a hundred mature oocysts of 50 um diameter could be observed at day 15 post-feed. Some of these mature oocysts had progressed into sporozoites but only a very few appeared upon disruption of the salivary glands.

To perform a genetic cross between *Pvs*EH and *Pvs*EL, a mixed inoculum containing equal proportions of *Pvs*EH and *Pvs*EL parasites was injected into CBA mice and a mosquito feed was performed on both day 3 and day 4 post-infection to increase the chances of a successful transmission (Additional 7 B). For each feed, around 160 female *A. stephensi* mosquitoes were allowed to take a blood meal from two anaesthetized mice at 24°C for 40 minutes without interruption.

Upon inspection of mosquito midguts for the presence of oocysts on day 9 post-feed, 100% infection was observed (all midguts inspected contained oocysts) for both day 3 and day 4 feeds. Around 25-100 oocysts were found per midgut in day 3 fed mosquitoes and 5-40 oocysts per midgut in day 4 fed mosquitoes. On day 12 post-feed, mature oocysts and also a high number of sporozoites were found in the midguts, but upon disrupting the salivary glands on day 20 post-feed, only a few sporozoites were found in the suspension.

Sporozoites from day 3 and 4 fed mosquitoes were injected into ICR mice (D3 and D4 respectively) and five days later, both mice became positive for blood stage parasites. In order to confirm that a genetic cross has taken place, four clones were obtained from D4 by limiting dilution to screen for presence of both *Pvs*EH and *Pvs*EL alleles within the chromosomes. Based on the SNPs identified between *Pvs*EH and *Pvs*EL, we amplified 600 to 1,000 bp regions from polymorphic genes on both ends of the 14 chromosomes that contained isolate-specific SNPs and performed Sanger sequencing of the amplicons (primer sequences in Additional file 6).

Both *Pvs*EL-specific (11) and *Pvs*EH-specific (17) markers were found in the 28 markers sequenced (one marker, PVSEL_0600390, could not be amplified). Also, five chromosomes clearly showed evidence of chromosomal cross-over since they contained markers from both isolates (see Figure 4A), thus confirming a successful *P. vinckei* genetic cross. However, all four clones had the same pattern of recombination which suggests that the diversity of recombinants in the cross- progeny was low and a single recombinant parasite might have undergone significant clonal expansion.

**Figure 4.**
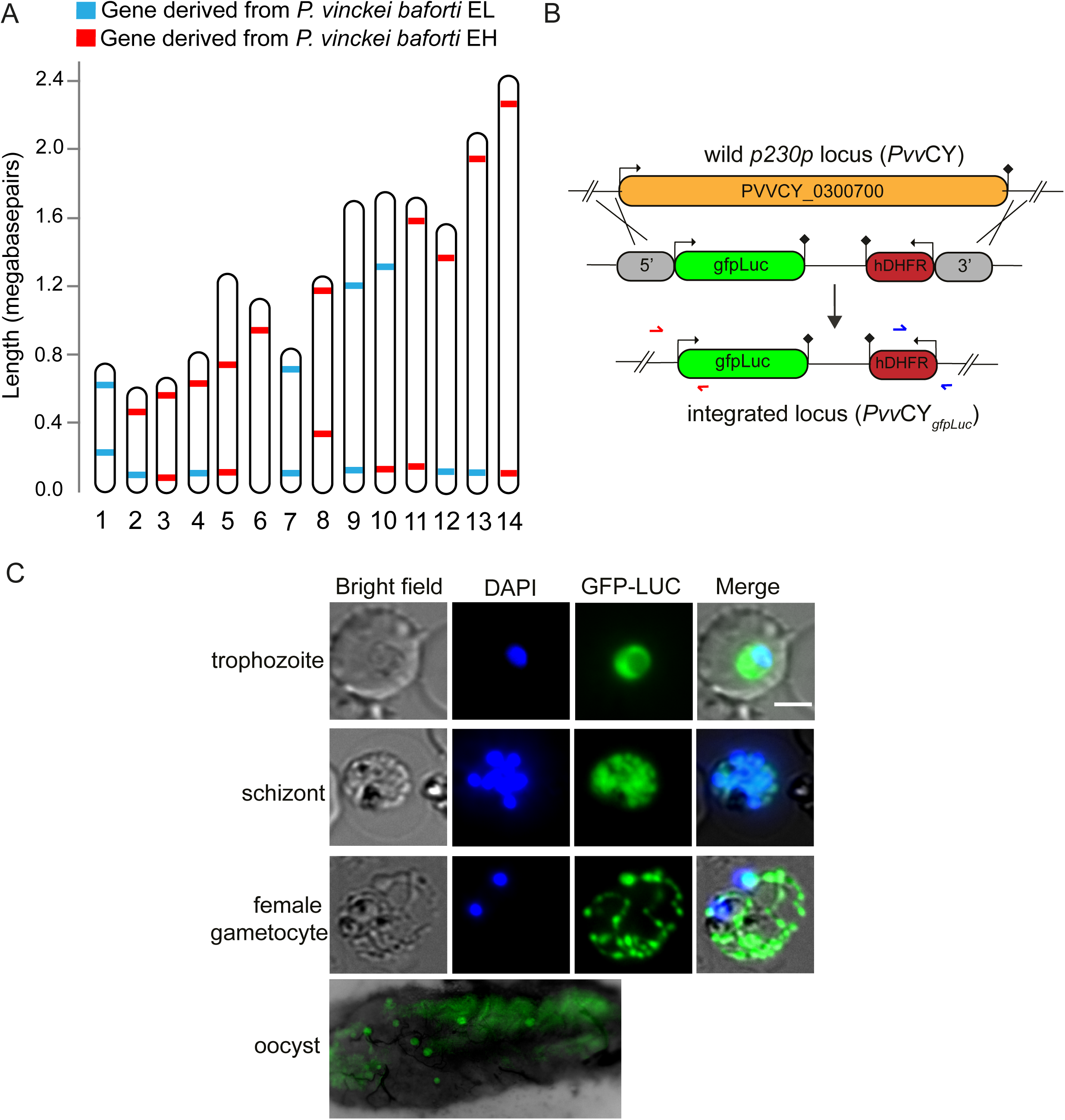
Phenotypic variation and genetics in *Plasmodium vinckei* parasites. A) Schematic of isolate-specific genetic markers detected in clonal line of *Pvs*EL X *Pvs*EH cross progeny by Sanger sequencing. Genetic markers from both EH (red) and EL (blue) isolates were detected in the crossed progeny proving successful genetic crossing. B) Schematic of homologous recombination-mediated insertion of a gfp-luciferase cassette into the p230p locus in *P. vinckei* CY. C) GFP expression in different blood stages of *Pvv*CY and luciferase expression of *Pvv*CY oocysts in mosquito midgut.

### *P. vinckei* parasites are amenable to genetic manipulation

We asked if *P. vinckei* parasites can be genetically modified by applying existing transfection and genetic modification techniques routinely used in other RMPs. *Plasmodium v. vinckei* CY was chosen to test this because the isolate naturally established a synchronous infection in mice and reaches a high parasitaemia, which results in an abundance of schizonts for transfection. We aimed to produce a *Pvv*CY line that constitutively expresses GFPLuc (green fluorescent protein-firefly luciferase) fusion protein, similar to those produced in *P. berghei* and *P. yoelii* [52, 53]. A recombination plasmid, *pPvvCY-* Δ*p230p-gfpLuc*, was constructed to target and replace the dispensable wildtype P230p locus in *P. v. vinckei* CY (PVVCY_0300700) with a gene cassette encoding for GFPLuc and a *hdhfr* selectable marker cassette (Figure 4B).

Transfection of purified *Pvv*CY schizonts with 20 µg of linearized p*Pvv*CY-Δp230p-gfpLuc plasmid by electroporation, followed by marker selection using pyrimethamine yielded pyrimethamine-resistant transfectant parasites (PvGFP-Luc_con_) on day 6 after drug treatment. Stable transfectants were cloned by limiting dilution and plasmid integration in these clones was confirmed by PCR. Constitutive expression of GFPLuc in PvGFP-Luc_con_ asexual and sexual blood stage parasites was confirmed by fluorescence live cell imaging (Figure 4C). GFPLuc expression in PvGFP-Luc_con_ oocysts was confirmed by fluorescence imaging of mosquito midguts 7 days after blood meal.

## Discussion

Of the four RMP species that have been adapted to laboratory mice, *P. berghei*, *P. yoelii* and *P. chabaudi* have been extensively used to investigate malaria parasite biology. Adopting these RMPs as tractable experimental models has been facilitated by continuous efforts in characterizing their phenotypes, sequencing their genomes and establishing protocols for parasite maintenance, genetic crossing and genetic modification. Here, we extend these efforts to *Plasmodium vinckei*.

We have systematically studied ten *P. vinckei* isolates and produced a comprehensive resource of their reference genomes, transcriptomes, genotypes and phenotypes to help establish *P. vinckei* as a useful additional experimental model for malaria.

Enzyme variation and molecular phylogeny studies indicate that the five subspecies of *P. vinckei* have diverged significantly from each probably due to the geographical isolation of these parasites in different locations around the African Congo basin. This diversity calls for a reference genome for each subspecies in order to capture large-scale changes in their genomes such as chromosomal structural variations and gene copy number variations that might have played a role in their subspeciation. To accurately capture these events, we used a combination of Pacbio and Illumina sequencing that allowed us to produce an end-to-end assembly of *P. vinckei* chromosomes. This, coupled with manual curation of the predicted gene models, led to the creation of five high-quality reference genomes for *P. vinckei* that are a significant improvement to the existing fragmented genomes available for *P. v. vinckei* and *P. v. petteri*.

Comparative synteny analysis between *P. vinckei* and other RMP genomes reveals structural variations at both the species and the subspecies levels. Assuming that the observed variations have occurred only once, a putative pathway of genome rearrangements during RMP evolution can be inferred. No rearrangements have occurred during *P. berghei* and *P. yoelii* speciation and their genomes are likely to be identical to the RMP ancestor [41]. A reciprocal translocation between chromosomes VIII and X has accompanied the speciation of *P. vinckei*, and this is mutually exclusive from the reciprocal translocation between chromosome VII and IX that has occurred during *P. chabaudi* speciation. Following this, there has been a small inversion in chromosome X during the subspeciation of *P. v. vinckei* and translocations between chromosomes V, VI and XIII during the subspeciation of *P. v. petteri* (which are then carried over to *P. v. baforti*).

We generated additional sequencing data for several *P. vinckei* isolates and made available at least two genotypes per *P. vinckei* subspecies (except for *P. v. vinckei* for which only one isolate is available) so as to facilitate future studies that might employ *P. vinckei* parasites to study phenotype-genotype relationships. Similarly, we also supplemented the existing genotype information for other RMPs by sequencing several isolates from additional subspecies of *P. chabaudi* and *P. yoelii*. Our data thus comprises of genotypes from sympatric species from each region of isolation allowing us to re-evaluate the genotypic diversity and evolution among RMP isolates.

A genome-wide SNP-based phylogeny shows that the divergences between different subspecies are proportional to the level of isolation of the habitat for all RMP species. *Plasmodium vinckei*, *P. yoelii* and *P. chabaudi* isolates from sites in Cameroon have very similar genotypes to their counterparts in the Central African Republic denoting similar evolutionary pressures and perhaps the presence of gene flow across these regions, while isolates from Brazzaville (Congo) are more diverged probably due to the different environmental conditions in these locations [40].

Subspecies from West Nigeria and the DRC are highly diverged compared to subspecies from the rest of Africa. The distinctiveness of *P. berghei* and *P. v. vinckei,* both from the DRC is most likely due to climactic and host-vector differences in the highland forests of Katanga. Highland forests are an altitude of 1000-7000 m with mean temperature of 21C whereas the lowland forests lie at an altitude less than 800m with a mean temperature of 25 C. Different host-vector systems are prevalent in the lowland forests (*Grammomys poensis (previously known as Thamnomys rutilans)* - specific mosquito species unknown) and the highland Katangan forests (*Grammomys surdaster* -*Anopheles dureni millecampsi*). The associated selection pressures seem to have mainly influenced their transmission, as reflected by their lower optimal transmission temperatures and the high Ka/Ks ratios observed for three proteins that play critical functions in this process. Recently, several more rodent host and mosquito vector species have been identified in the forests of Gabon [54] implying that a diverse set of host-vector systems could have existed for RMPs. Thus, diversification of RMP species into several subspecies within these isolated ecological niches might have been driven by evolutionary forces resulting from the diverse host, vector and environmental conditions experienced at each locale.

Malaria parasite genomes contain several highly polymorphic multigene families located in the sub-telomeric chromosomal regions that encode a variety of exported proteins involved in processes such as immune evasion, cytoadherence, nutrient uptake and membrane synthesis. Multigene families are thought to have evolved rapidly under the influence of immune and other evolutionary pressures resulting in copy number variations and rampant sequence reshuffling that ultimately leads to phenotypic plasticity in *Plasmodium*.

Previously, phylogenetic analyses of *pir, fam-a and fam-b* genes from three RMP species have shown that structurally distinct genes exist within these families forming robust clades with varied levels of orthology/paralogy. Identifying sub-families that have structurally diversified within the multigene families can help to better understand their functions and to this end, we constructed phylogenetic trees for the ten multigene families with genes from all four RMP species. Due to the scale of the analysis, we applied automated trimming to our alignments and limited our tree inference method to maximum likelihood. While this resulted in poor bootstrap values for some clades in the *pir*, *fam-a* and *fam-b* trees compared to previous phylogenetic analyses [25, 26], our method was able to retrieve similar tree topologies to those previously inferred and in general produced trees with good nodal support for the rest of the multigene families.

Robust pan-RMP clades identified in our study represent ancestral lineages consisting of structural orthologs that perform conserved functions across all RMPs and will be useful for future work with these families. We show that certain ancestral lineages can expand in a particular species or subspecies in response to selective pressures resulting in distinct evolutionary histories for each family. For example, the *pir* family expansion is mostly species-specific and driven by frequent gene conversion after speciation, whereas the expansion of the *fam-a* gene repertoire seems to have occurred initially in the RMP ancestor followed by species-specific expansions.

Inclusion of *P. vinckei pirs*in the RMP *pir* family phylogeny show that *P. vinckei pir*s do not form independent clades of their own and instead populate three *P. chabaudi*-dominant clades. This suggests that some of the *pir* clades were established earlier on when the classical *vinckei* and *berghei* group of parasites split from their common RMP ancestor resulting in *vinckei* group-specific clades like L1, S7 and S1g.

Similarly, the addition of *P. vinckei* genes resolved the ancestral clade of internal *fam-a* genes into several well-supported *vinckei* group clades and a *berghei* group clade. We observe similarly high level of orthology between *P. chabaudi* and *P. vinckei* genes in other multigene families forming several *vinckei* group clades in contrast to more species-specific clades of paralogous genes in *P. berghei* and *P. yoelii*. For example, within the *fam-d* family, five ancestral lineages can be identified in the *vinckei* group as opposed to only one paralogous *P. yoelii*-specific expansion within the *berghei* group. Taken together, it seems that family expansions in *P. chabaudi* and *P. vinckei* have occurred in the common *vinckei* group ancestor prior to speciation and that multigene families have evolved quite differently across the *vinckei* and *berghei* groups of RMPs. These might be related to the striking differences in the basic phenotypes of these two groups of parasites.

We also observed size expansions in the *ema1* and *fam-c* families within the non-Katangan *P. vinckei* parasites, all being isolates from the lowlands around the Congo Basin. *Ema1* family expansions seem to be specific to lowlands dwelling *vinckei* group parasites as they are expanded in both non-Katangan *P. vinckei* and *P. chabaudi.* However, unlike *P. chabaudi*, the duplicated gene members in non-Katangan *P. vinckei* are all pseudogenized by a S5X mutation effectively rendering the functional repertoire to be just 6-8 genes, similar to highlands dwelling Katangan *P. v. vinckei*. Thus, it could be speculated that even under similar selective pressures, *ema1* family expansions contribute to parasite fitness in *P. chabaudi* but may not be required for the survival of sympatric *P. vinckei* parasites. The *P. vinckei ema1* pseudogenes could still serve as silent donor genes that recombine into functional variants to bring about antigenic variation [55]. In the case of the *fam-c* gene family, the expansion is specific to non-Katangan *P. vinckei* subspecies since *P. chabaudi*, *P. yoelii* and *P. v. vinckei* all have similar repertoire sizes. The expansions seem to be driven by gene duplications initially in their non-Katangan common ancestor and again after subspeciation.

The effect of the difference in habitats is even more pronounced in the Katangan parasite, *P. v. vinckei*. It has a smaller genome and a compact multigene family repertoire reminiscent of the only other Katangan isolate, *P. berghei* and its genetic distance from other members of the *P. vinckei* clade is in the same order of magnitude as that between separate species within the RMPs. The reduced multigene family repertoire mainly consists of members belonging to pan-RMP or *vinckei*-group specific ancestral lineages making it an ideal *vinckei* group parasite to study the localization and function of variant proteins.

We tested whether *Pvv*CY was amenable to genetic manipulation using standard transfection protocols already established for other RMPs. We were able to successfully knock-in a GFP-luciferase fusion cassette to *Pvv*CY to produce a GFP-Luc reporter line for *P. v. vinckei,* following a transfection protocol routinely used for modifying *P. yoelii* in our lab [44, 56]. We were able to visualise GFP-positive parasites during different blood stages and in oocysts thus confirming stable GFPLuc expression. We were unable to visualise other life stages (sporozoites and liver stages) due to our failure to produce viable salivary gland sporozoites in this parasite.

The transfection of *P. chabaudi* has been challenging due to its slow proliferation rate and schizont sequestration resulting in low merozoite yield, thus necessitating optimized transfection protocols. In contrast, *Plasmodium v. vinckei* reaches high parasitaemia without being immediately lethal to the host (90% parasitaemia on day 6) and is highly synchronous yielding a large number of schizonts. A predominant population of schizonts appear near midnight in *P. v. vinckei* infections, at which point, they can be Percoll-purified from exsanguinated blood and transfected with DNA.

*P. vinckei* and *P. chabaudi*, while being distinct species, share several characteristics that are common among *vinckei* group RMPs, such as a predilection for mature erythrocytes, synchronous infections and the sequestration of schizonts from peripheral circulation [33, 37, 57–59]. Thus, *P. v. vinckei* can serve as an ideal experimental model for functional studies targeting these aspects of parasite biology.

The availability of several RMP isolates with phenotypic differences aids their use in study of parasite fitness and transmission success in mixed infections [60, 61] and for the identification of genes involved in parasite virulence, strain-specific immunity, drug resistance and host-cell preference using genetic crosses [44, 62–64]. With this in mind, we studied the virulence of ten *P. vinckei* isolates to identify differences in their growth rate and their effect on the host.

Some of these isolates have been previously characterized [40], but we systematically profiled additional representative isolates for each subspecies (where available) under comparative conditions in the same host strain. We identified pairs of isolates with contrasting virulence phenotypes within two *P. vinckei* subspecies – *P. v. petteri (Pvp*CR and *Pvp*BS) and *P. v. baforti* (*Pvs*EH and *Pvs*EL or *Pvs*EE). These isolate pairs would be ideal candidates for studies utilising genetic crossing to identify genetic *loci* linked to virulence using Linkage Group Selection [44].

Since *P. vinckei* subspecies have significantly diverged from each other, isolates within the same subspecies are more likely to recombine than isolates from different subspecies. However, intra-specific hybrids between *P. v. petteri* and *P. v. baforti* may also be possible (as demonstrated earlier in *P. yoelii* [65]) since these two subspecies are closely related (see Figure 2C). However, difficulties in transmitting *P. vinckei* parasites have been reported previously with either the gametocytes failing to produce midgut infections or sporozoites failing to invade the salivary glands or infections resulting in non-infective sporozoites [27, 30, 35]. Repeated attempts to create a cross between two *P. vinckei baforti* isolates failed to produce any detectable recombinants due to low frequency of mosquito transmission [35]. Here, we renewed these efforts with different *P. vinckei* isolates to see if we could establish a *P. vinckei* genetic cross. Two attempts were made to create a *pvp*CR X *pvp*BS cross and further two attempts were made to create a *pvs*EL X *pvs*EH cross. However, in all attempts the sporozoites failed to optimally invade the salivary glands and we managed to isolate only a few in the *P. v. subsp* cross, subsequently obtaining a cross progeny in mice. While we were able to demonstrate a successful genetic cross by showing the presence of alleles from both isolates in the cross progeny, the recombinant diversity was quite low probably due to the transmission bottleneck. We are currently further investigating the optimal conditions for transmitting *P. vinckei*.

## Conclusions

In this study, we have created a comprehensive resource for the rodent malaria parasite *Plasmodium vinckei*, comprising of five high-quality reference genomes, and blood stage-specific transcriptomes, genotypes and phenotypes for ten isolates. We have employed state-of-the-art sequencing technologies to produce largely complete genome assemblies and highly accurate gene models that were manually polished based on strand-specific RNA sequencing data. The unfragmented nature of our genome assemblies allowed us to characterize structural variations within *P. vinckei* subspecies, which, to the best of our knowledge, is the first time that large-scale genome re-arrangements have been found among subspecies of a *Plasmodium* species.

The biological or phenotypic significance, if any, of such alterations are poorly understood, but it seems likely that they may drive speciation through the promotion of reproductive isolation of species or subspecies. Through our extensive sequencing efforts, we have generated genotype data for seventeen RMP isolates comprising of five *P. vinckei*, four *P. yoelii* and one *P. chabaudi* subspecies, thus making at least one genotype available for all subspecies of the RMP that previously lacked any sequencing data. We also systematically characterised the virulence phenotypes of the ten *P. vinckei* isolates to capture the phenotypic diversity among them. Combined, these efforts will greatly aid genetic linkage studies to resolve genotype-phenotype relationships.

In order to understand the evolutionary relationships among the RMP isolates, we have carried out a combination of analyses to describe the genotypic diversity molecular evolution of these parasites. While our phylogenies more or less agree with previous biochemical and molecular data-based studies, our reconstruction based on sequence variations on a genome scale provides higher resolution to the divergence estimates. Taking advantage of the high-quality RMP genomes produced from our work and previous studies, we also undertook a comprehensive phylogenetic analysis of multigene families across all RMP species and identify various structurally diversified sub-families with distinct evolutionary histories. This will enable future studies on the critical role of multigene families in parasite adaptation, and to aid this, we have made searchable and interactive versions of the phylogenies publicly available through the iTOL online tool [66].

While genome rearrangements have occurred during speciation and sub speciation events, diversification of the multigene families seem to have occurred earlier when the RMPs split into *vinckei* and *berghei* groups of parasites. Thus, structural, copy number and nucleotide-level variations among the RMPs have occurred at various points during the evolution of RMPs in response to a variety of evolutionary pressures. The gene expression data from our study, covering specific blood stages for some *P. vinckei* subspecies, show conserved expression of multigene family members across RMPs. While not comprehensive, it complements existing RMP transcriptomes and will aid functional studies in the *P. vinckei* model. Taken together, our study provides a comprehensive view of the phenotypic and genotypic diversity within RMPs and functional diversification of the multigene families in response to selection pressures.

The synchronicity of *P. v. vinckei* infection and its unique ability to sustain high parasitaemia without killing its host culminating in good schizont yields make this parasite an attractive model for reverse genetics studies, especially those on multigene families owing to its reduced repertoire. We have successfully demonstrated genetic manipulation in *P. v. vinckei* but encountered difficulties in producing large numbers of recombinant parasites through genetic crossing. Attempts to transmit isolates from three different *P. vinckei* subspecies in *A. stephensi* mosquitoes failed in our hands as sporozoites repeatedly failed to infect the salivary glands. Careful optimisation of transmission parameters and serial mosquito passages of the *P. vinckei* parasites might help in improving their transmission efficiency and could aid genetic linkage studies with these parasites.

## Methods

### Parasite lines and experiments using mice and mosquitos

The parasite lines used in this study and their original isolate information are detailed in Supplementary Table 1. Frozen parasite stabilates of cloned or uncloned lines were revived and inoculated intravenously into ICR mice. Five *P. vinckei* isolates (*Pvv*CY, *Pvb*DA, *Pvb*DB, *Pvl*DE and *Pvs*EE) and the *P. yoelii nigeriensis* isolate (*Pyn*D) were uncloned stabilates and were cloned by limiting dilution to obtain clonal parasite lines.

Laboratory animal experimentation was performed in strict accordance with the Japanese Humane Treatment and Management of Animals Law (Law No. 105 dated 19 October 1973 modified on 2 June 2006), and the Regulation on Animal Experimentation at Nagasaki University, Japan. The protocol was approved by the Institutional Animal Research Committee of Nagasaki University (permit: 12072610052).

Six to eight weeks old female ICR or CBA mice were used in all the experiments.

The mice were housed at 23°C and maintained on a diet of mouse feed and water. Mice infected with malaria parasites were given 0.05% para-aminobenzoic acid (PABA)- supplemented water to assist parasite growth.

All mosquito transmission experiments were performed using *Anopheles stephensi* mosquitoes were housed in a temperature and humidity-controlled insectary at 24°C and 70% humidity. Mosquito larvae were fed with mouse feed and yeast mixture and adult mosquitoes were maintained on 10% glucose solution supplemented with 0.05% PABA.

### Parasite growth profiling

For each isolate, an inoculum containing 1 X 10^6^ parasitized RBCs was injected intravenously to five CBA mice. Blood smears, haematocrit readings (Beckman Coulter Counter) and body weight readings were taken daily for 20 days or until host mortality to monitor parasitaemia, anaemia and weight loss. Blood smears were fixed with 100% methanol and stained with Geimsa’s solution. The average parasitaemia was calculated from parasite and total RBC counts taken at three independent microscopic fields.

### Genomic DNA isolation and whole genome sequencing

Parasitized whole blood was collected from the brachial arteries of infected mice and blood sera was removed by centrifugation. RBC pellets were washed once with PBS and leukocyte-depleted using CF11 (Sigma Cat# C6288) cellulose columns. Parasite pellets were obtained by gentle lysis of RBCs with 0.15% saponin solution. Genomic DNA extraction from the parasite pellet was performed using DNAzol reagent (Invitrogen CAT # 10503027) as per manufacturer’s instructions.

Single-molecule sequencing was performed for five *P. vinckei* isolates. 5-10 ug of gDNA was sheared using a Covaris g-TUBE shearing device to obtain target sizes of 20kB (for *Pvv*CY, *Pvb*DA and *Pvp*CR) and 10kB (for samples *Pvl*DE and *Pvs*EL). Sheared DNA was concentrated using AMPure magnetic beads and SMRTbell template libraries were generated as per Pacific Biosciences instructions. Libraries were sequenced using P6 polymerase and chemistry version 4 (P6C4) on 3-6 SMRT cells and sequenced on a PacBio RS II. Reads were filtered using SMRT portal v2.2 with default parameters. Read yields were 352,693, 356,960, 765,596, 386,746 and 675,879 reads for *Pvv*CY, *Pvb*DA, *Pvl*DE, *Pvp*CR and *Pvs*EL respectively totalling around 2.7 to 4.7 Gb per sample. Mean subread lengths ranged from 6.15 to 9.1 kB. N50 of 11.7 kB and 19.2 kB were obtained for 10 and 20 kB libraries respectively.

PCR-free Illumina sequencing was performed for all RMP isolates. 1-2 ug of DNA was sheared using Covaris E series to obtain fragment sizes of 350 and 550bp. 350bp and 550bp PCR-free libraries were prepared using TruSeq PCR-free DNA library preparation kits according to the manufacturer’s instructions. Libraries were sequenced on the Illumina HiSeq2000 platform with 2 × 100bp paired-end read chemistry. Read yields ranged from 8-22 million reads for each library (see Additional File 1).

### Genome assembly and annotation

Genome assembly from long single molecule sequencing reads was performed using FALCON (v0.2.1)[67] with length cutoff for seed reads used for initial mapping set as 2,000bp and for pre-assembly set as 12,000bp. The falcon sense options were set as- ”–min idt 0.70 –min cov 4 –local match count threshold 2 –max n read 200” and overlap filtering settings were set as ”–max diff 240 –max cov 360 –min cov 5 –bestn 10”. 28-40 unitigs were obtained and smaller unitigs were discarded as they were exact copies of the regions already present in the larger unitigs.

PCR-free reads were used to correct base call errors in the unitigs using ICORN2 [68], run with default settings and for 15 iterations. The unitigs were classified as chromosomes based on their homology with *P. chabaudi* chromosomes (GeneDB version 3). In *Pvl*DE and *Pvs*EL samples, some of the chromosomes were made of two to three unitigs with overlapping ends which were then fused and the gaps were removed manually. Apicoplast and mitochondrial genomes were assembled from PCR-free reads alone using Velvet assembler [69].

Syntenic regions between genome sequences were identified using MUMmer v3.2 [70]. Synteny breakpoints were identified manually and were confirmed not to be misassemblies by verifying that they had continuous read coverage from PacBio and Illumina reads. Artemis Comparison tool [71, 72] and Integrative Genomics Viewer [73] were used for this purpose. The structural variations were illustrated using CIRCOS [74].

*De novo* gene predictions were made using AUGUSTUS [75] trained on *P. chabaudi* gene models. RNA sequencing reads were mapped onto the reference genome using TopHat [76] to infer splice junctions. AUGUSTUS predicted gene models, junctions.bed file from TopHat and *P. chabaudi* gene models were fed into MAKER [77] to create consensus gene models that were then manually curated based on RNAseq evidence in Artemis Viewer and Artemis Comparison tool [71, 72]. Ribosomal RNA (rRNA) and transfer RNA (tRNA) were annotated using RNAmmer v1.2 [78]. Gene product calls were assigned to *P. vinckei* gene models based on above identified orthologous groups using custom scripts. Functional domain annotations were inferred from InterPro database using InterProScan v5.17 [79]. Transmembrane domains were predicted by TMHMMv2.0 [80], signal peptide cleavage sites by SignalP v4.0 [81], presence of PEXEL/VTS motif detected using ExportPredv4.0 [82] (with PEXEL score cutoff of 4.3).

### Transcriptomics

Total RNA was isolated for four *P. vinckei* isolates (*Pvb*DA, *Pvp*CR, *Pvl*DS and *Pvs*EL) from mixed blood stages using TRIzol (Invitrogen) following the manufacturer’s protocol. For *Pvp*CR and *Pvl*DS, additionally, total RNA was isolated from ring, trophozoite and gametocyte enriched fractions obtained using a Nycodenz gradient.

Strand-specific mRNA sequencing was performed from total RNA using TruSeq Stranded mRNA Sample Prep Kit LT (Illumina) according to the manufacturer’s instructions. Briefly, polyA+ mRNA was purified from total RNA using oligo-dT dynabead selection. First strand cDNA was synthesised using randomly primed oligos followed by second strand synthesis where dUTPs were incorporated to achieve strand-specificity. The cDNA was adapter-ligated and the libraries amplified by PCR. Libraries were sequenced in Illumina Hiseq2000 with paired-end 100bp read chemistry.

Stage-specific RNAseq data for *Pvv*CY’s intraerythrocytic growth stages were obtained from an earlier study [50]. Gene expression was captured every 6 hours during *Pvv*CY’s 24 h IDC with three replicates, of which 6h, 12h and 24h timepoints were used in this study to denote gene expression at ring, trophozoite and schizont stages respectively. Similarly, for *P. chabaudi* AS, gene expression was captured every 3h during its IDC with two replicates in a recent study [56], of which the 5.5h, 11.5h and 23.5 h timepoints on day 2 were chosen to denote ring, trophozoite and schizont stages respectively. *P. yoelii* and *P. berghei* transcriptome data were obtained from [26] and [44] respectively.

### SNP calling and molecular evolution analysis

Illumina paired-end reads for a total of 30 RMP isolates produced in this study or sourced from previous studies (see Additional File 13) were used for SNP calling. In the case of isolates sequenced in this study, the 350bp fragment size PCR-free sequencing data was used. First, to produce a high quality pan-RMP SNP dataset for phylogeny construction, all quality-trimmed reads were mapped onto the *Pvv*CY reference genome using BWA tool [83] with default parameters. MAPQ values of the mapped reads were fixed and duplicated reads removed using CleanSam, FixMateInformation and MarkDuplicates commands in picardtools (http://broadinstitute.github.io/picard) and only uniquely mapped reads were retained using samtools with parameter -q 1 (http://www.htslib.org/). Raw SNPs were called from the mapped reads using samtools mpileup and bcftools with following parameters-minimum base quality of 20, minimum mapping quality of 10 and ploidy of 1. SNPs with quality (QUAL) less than 20, read depth (DP) less than 10, mapping quality (MQ) less than 2 and allele frequency (AF1) less than 80% were removed. Further, only SNPs present in protein-coding genes were retained and those present in low-complexity regions (predicted by DustMasker [84]) and sub-telomeric multigene family members were excluded.

The filtered SNPs from different samples were merged and SNP positions with missing calls in more than six samples were removed. This filtered high-quality set of 1,020,956 SNP positions were used to infer maximum likelihood phylogeny (see Additional File 7).

For inferring Ka/Ks ratios between *P. vinckei* isolates and *Pvv*CY, filtered SNPs obtained above were merged as before but excluding *Pvp*BS due to its high missing call rate. Only SNP positions with no missing calls in any sample were retained and morphed onto *Pvv*CY gene sequences using gatk command FastaAlternateReferenceMaker [85] to produce isolate-specific gene sequences which were then used for pairwise sequence comparisons to identify synonymous and non-synonymous substitutions. Ka/Ks ratios were calculated using KaKs Calculator [86] and averaged across isolates if more than one was available for a subspecies.

For comparisons against *P. v. petteri*, *P. yoelii* and *P. chabaudi*, sample reads were mapped onto *Pvp*CR, *P. yoelii* 17X and *P. chabaudi* AS genomes respectively and subsequent steps were followed as before. Similar to *Pvp*BS, *Pys*EL was excluded from Ka/Ks analysis due to high missing rate.

### Phylogenetic analysis

For constructing species-level phylogenies, orthologous proteins were identified between the five *P. vinckei* genomes, three RMP genomes, *P. falciparum, P. knowlesi* and *P. vivax* genomes using OrthoMCL v2.0.9 [87] with inflation parameter as 1.5, BLAST hit evalue cutoff as 1e^-5^ and percentage match cutoff as 50%.

One-to-one orthologous proteins from each of the 3,920 ortholog groups that form the core proteome were aligned using MUSCLE [88]. Alignments were trimmed using trimAl [89] removing all gaps and concatenated into a partitioned alignment using catsequence. An initial RAxML [90] run was performed on individual alignments to identify best amino acid substitution model under the Akaike Information Criterion (--auto-prot=aic). These models were then used to run a partitioned RAxML analysis on the concatenated protein alignment using PROTGAMMA model for rate heterogeneity.

For constructing isolate-level phylogeny, the vcf files containing high-quality SNPs were first converted to a matrix for phylogenetic analysis using vcf2phylip (https://github.com/edgardomortiz/vcf2phylip). RAxML tree inference was performed using GTRGAMMA model for rate heterogeneity along with ascertainment bias correction (--asc-corr=stamatakis) since we used only variant sites.

Maximum likelihood trees for multigene families were constructed based on nucleotide sequence alignments of member genes that included intron sequences if present (except in the case of *pir* family where introns were excluded). Alignments were performed using MUSCLE with default parameters, frame-shifts edited manually in AliView [91] followed by automated trimming with trimAl using -gappyout parameter. In all the phylogenies, bootstrapping was conducted until the majority-rule consensus tree criterion (-I autoMRE) was satisfied (usually 150-300 replicates). Phylogenetic trees were visualized and annotated in the iTOL server [66].

### Plasmid construction and transfection in *P. vinckei*

The p*Pvv*CY-p230p-gfpLuc plasmid was constructed using MultiSite Gateway cloning system (Invitrogen). attB-flanked 5’and 3’homology arms were obtained by amplifying 800bp regions upstream and downstream of PVVCY_0300700. These fragments were subjected to independent BP recombination with pDONRP4-P1R (Invitrogen) to generate entry plasmids pENT12-5U and pENT41-3U, respectively. Similarly, the gfpLuc cassette from pL1063 was amplified and subjected to LR reaction to obtain pENT23-gfpLuc. BP reaction was performed using the BP Clonase II enzyme mix (Invitrogen) according to the manufacturer’s instructions.

*Plasmodium vinckei vinckei* CY schizont-enriched fraction was collected by differential centrifugation on 50% Nycodenz in incomplete RPMI1640 medium, and 20 ug of ApaI- and StuI-double digested linearized transfection constructs were electroporated to 1 x 10^7^ of enriched schizonts using a Nucleofector device (Amaxa) with human T-cell solution under program U-33. Transfected parasites were intravenously injected into 7-week-old ICR female mice, which were treated by administering pyrimethamine in the drinking water (0.07 mg/mL) 24 hours later for a period of 4-7 days. Drug resistant parasites were cloned by limiting dilution with an inoculum of 0.3 parasites/100 uL injected into 10 female ICR mice. Two clones were obtained, and integration of the transfection constructs was confirmed by PCR amplification with a unique set of primers for the modified p230p gene locus. Live imaging of parasites was performed on thin smears of parasite-infected blood prepared on glass slides stained with Hoechst 33342. Fluorescent and differential interference contrast (DIC) images were captured using an AxioCam MRm CCD camera (Carl Zeiss, Germany) fixed to an Axio imager Z2 fluorescent microscope with a Plan-Apochromat 100 ×/1.4 oil immersion lens (Carl Zeiss) and Axiovision software (Carl Zeiss). GFP-expressing *P. vinckei* oocysts in mosquito midguts were imaged in SMZ25 microscope (Nikon).

### Mosquito transmission and genetic crossing of *P. vinckei* parasites

To determine the optimal transmission temperature for *P. vinckei baforti* isolates, infected CBA mice were anaesthetized on day 3 post-inoculation and ∼100 female *Anopheles stephensi* mosquitoes (7 to 12 days post emergence) were allowed to take a blood meal for 30 min without interruption after confirming presence of gametocytes by microscopy. Three batches of ∼100 mosquitoes were fed at three different temperatures - 21°C, 23°C and 26°C. The fed mosquitoes were maintained at the feed temperatures and at 70% humidity. To check for presence of oocysts/sporozoites, mosquitoes were dissected, and their midguts or salivary glands were suspended in a drop of PBS solution atop a glass slide, covered by a coverslip and studied under a microscope.

For genetic crossing, isolates were harvested from donor mice and mixed to achieve a 1:1 ratio and 1 x 10^6^ parasites of this mixture was inoculated into four female CBA mice. Three days after inoculation, after confirming the presence of gametocytes, two infected CBA mice were anaesthetized and placed on two mosquito cages, each containing around 80 mosquitoes each. Mosquitoes were allowed to feed on the mice without interruption for 40 minutes at 24°C. A fresh feed was again performed on the 4th day post-inoculation with the other two CBA mice and two fresh cages of mosquitoes. 5-10 female mosquitoes from each cage were dissected on the 9th and 12th day after the blood meal to check for presence of oocysts in the mosquito midguts. Twenty days after the blood meal, the mosquitoes were dissected and the salivary glands were removed, placed in 0.5-0.7 ml PBS solution and gently disrupted to release sporozoites. The suspensions from day 3 and day 4 feeds were injected intravenously into an ICR mouse each. When the mice were positive for blood-stage parasites, they were sub-inoculated into ten ICR mice with an inoculum of 0.6 parasites/100uL to obtain clones from the potential cross progeny by limiting dilution. Eight days post infection, four mice were positive for parasites and these clones were screened for the presence of both *Pvs*EH and *Pvs*EL alleles within the chromosomes.

## Supporting information

AdditionalFile1

AdditionalFile2

AdditionalFile3

AdditionalFile4

AdditionalFile5

AdditionalFile6

AdditionalFile7

AdditionalFile8

AdditionalFile9

AdditionalFile10

AdditionalFile11

AdditionalFile12

## Declarations

### Consent for publication

Not applicable

### Availability of data and materials

All genome sequences, gene annotations and sequencing data files generated in this study can be found in ENA Study: PRJEB19355. All the datasets would also be available via EuPathDB portal at the time of publication. All parasite resources will be made available to the scientific community via the BEI Resources (https://www.beiresources.org/). Searchable and interactive versions of the phylogeny trees produced in this study can be accessed at https://itol.embl.de/shared/2lCr6w0mdDENs.

### Authors’ contributions

RC and AP conceived the study. AR and RC conducted all rodent and mosquito experiments. AR and OG prepared sequencing libraries. AR, SK and RC conducted genetic cross experiments. AR collected data and performed all bioinformatic analyses. AR wrote the manuscript and all authors contributed to it.

## Acknowledgements

The authors would like to acknowledge the personnel at the Bioscience Core Laboratory (BCL) at King Abdullah University of Science and Technology for their help with next generation sequencing. We acknowledge the contribution of Professor Richard Carter, whose support of this work was crucial to its inception and completion.

## Authors’ information (optional)

Not applicable.

## Additional files

Additional file 1. Summary of rodent malaria parasite isolates used and DNA and RNA sequencing performed in this study.

Additional file 2. Infection profiles of ten *Plasmodium vinckei* isolates. Changes in parasitaemia, host RBC density and host weight during *P. vinckei* infections. Error bars show standard deviation of the readings within five biological replicates. † denotes host mortality.

Additional file 3. Daily readings of parasitaemia, host RBC density and host weight during infections of *Plasmodium vinckei* isolates.

Additional file 4. Assembly statistics of *Plasmodium vinckei* mitochondrial and apicoplast genomes.

Additional file 5. Chromosomal synteny breakpoints among *Plasmodium vinckei* genomes.

Additional file 6. 3,920 one-to-one orthologous group used for genome-wide protein alignment-based phylogeny.

Additional file 7. Single nucleotide polymorphisms among RMP isolates and Ka/Ks ratios for various pair-wise comparisons of homologous protein-coding genes.

Additional file 8. Copy number variations within multigene families and phylogenetic clade members.

Additional file 9. Maximum Likelihood trees for ten RMP multigene families.

Additional file 10. Gene-wise RNA-seq FPKM values for *Plasmodium vinckei petteri* CR, *P. v. lentum* DS (Rings, trophozoites and gametocyte stages), *P.v. brucechwatti* DA and *P. v. baforti* EL (mixed blood stages).

Additional file 11. Mosquito transmission and genetic cross experiments. Additional file 12. A) Circos figure showing rearrangements among four RMP species B) Gene alignment of pseudogenised ema1 genes.

## References

1. Carlton JM, Hayton K, Cravo PV, Walliker D Of mice and malaria mutants: unravelling the genetics of drug resistance using rodent malaria models. Trends Parasitol 2001, 17(5):236–242.

2. Culleton RL, Abkallo HM Malaria parasite genetics: doing something useful. Parasitol Int 2015, 64(3):244–253.

3. Matz JM, Kooij TW Towards genome-wide experimental genetics in the in vivo malaria model parasite Plasmodium berghei. Pathog Glob Health 2015, 109(2):46–60.

4. De Niz M, Heussler VT Rodent malaria models: insights into human disease and parasite biology. Curr Opin Microbiol 2018, 46:93–101.

5. Carter R Studies on enzyme variation in the murine malaria parasites Plasmodium berghei, P. yoelii, P. vinckei and P. chabaudi by starch gel electrophoresis. Parasitology 1978, 76(3):241–267.

6. Perkins SL, Sarkar IN, Carter R The phylogeny of rodent malaria parasites: simultaneous analysis across three genomes. Infect Genet Evol 2007, 7(1):74–83.

7. Ramiro RS, Reece SE, Obbard DJ Molecular evolution and phylogenetics of rodent malaria parasites. BMC Evol Biol 2012, 12:219.

8. Cravo PV, Carlton JM, Hunt P, Bisoni L, Padua RA, Walliker D Genetics of mefloquine resistance in the rodent malaria parasite Plasmodium chabaudi. Antimicrob Agents Chemother 2003, 47(2):709–718.

9. Langhorne J, Quin SJ, Sanni LA Mouse models of blood-stage malaria infections: immune responses and cytokines involved in protection and pathology. Chem Immunol 2002, 80:204–228.

10. Spence PJ, Jarra W, Levy P, Reid AJ, Chappell L, Brugat T, Sanders M, Berriman M, Langhorne J Vector transmission regulates immune control of Plasmodium virulence. Nature 2013, 498(7453):228–231.

11. Brugat T, Reid AJ, Lin J, Cunningham D, Tumwine I, Kushinga G, McLaughlin S, Spence P, Bohme U, Sanders M, et al Antibody-independent mechanisms regulate the establishment of chronic Plasmodium infection. Nat Microbiol 2017, 2:16276.

12. Stephens R, Culleton RL, Lamb TJ The contribution of Plasmodium chabaudi to our understanding of malaria. Trends Parasitol 2012, 28(2):73–82.

13. Prudencio M, Mota MM, Mendes AM A toolbox to study liver stage malaria. Trends Parasitol 2011, 27(12):565–574.

14. Guttery DS, Roques M, Holder AA, Tewari R Commit and Transmit: Molecular Players in Plasmodium Sexual Development and Zygote Differentiation. Trends Parasitol 2015, 31(12):676–685.

15. Pfander C, Anar B, Schwach F, Otto TD, Brochet M, Volkmann K, Quail MA, Pain A, Rosen B, Skarnes W et al A scalable pipeline for highly effective genetic modification of a malaria parasite. Nat Methods 2011, 8(12):1078–1082.

16. Marr E, Milne R, Anar B, Girling G, Schwach F, Mooney J, Nahrendorf W, Spence P, Cunningham D, Baker D et al An enhanced toolkit for the generation of knockout and marker-free fluorescent Plasmodium chabaudi [version 1; peer review: 2 approved]. Wellcome Open Research 2020, 5(71).

17. Janse CJ, Ramesar J, Waters AP High-efficiency transfection and drug selection of genetically transformed blood stages of the rodent malaria parasite Plasmodium berghei. Nat Protoc 2006, 1(1):346–356.

18. Jongco AM, Ting LM, Thathy V, Mota MM, Kim K Improved transfection and new selectable markers for the rodent malaria parasite Plasmodium yoelii. Mol Biochem Parasitol 2006, 146(2):242–250.

19. Bushell E, Gomes AR, Sanderson T, Anar B, Girling G, Herd C, Metcalf T, Modrzynska K, Schwach F, Martin RE et al Functional Profiling of a Plasmodium Genome Reveals an Abundance of Essential Genes. Cell 2017, 170(2):260–272 e268.

20. Stanway RR, Bushell E, Chiappino-Pepe A, Roques M, Sanderson T, Franke-Fayard B, Caldelari R, Golomingi M, Nyonda M, Pandey V et al Genome-Scale Identification of Essential Metabolic Processes for Targeting the Plasmodium Liver Stage. Cell 2019, 179(5):1112–1128 e1126.

21. Antonova-Koch Y, Meister S, Abraham M, Luth MR, Ottilie S, Lukens AK, Sakata-Kato T, Vanaerschot M, Owen E, Jado JC et al Open-source discovery of chemical leads for next-generation chemoprotective antimalarials. Science 2018, 362(6419).

22. de Koning-Ward TF, Gilson PR, Crabb BS Advances in molecular genetic systems in malaria. Nat Rev Microbiol 2015, 13(6):373–387.

23. Carlton JM, Angiuoli SV, Suh BB, Kooij TW, Pertea M, Silva JC, Ermolaeva MD, Allen JE, Selengut JD, Koo HL et al Genome sequence and comparative analysis of the model rodent malaria parasite Plasmodium yoelii yoelii. Nature 2002, 419(6906):512–519.

24. Hall N, Karras M, Raine JD, Carlton JM, Kooij TW, Berriman M, Florens L, Janssen CS, Pain A, Christophides GK et al A comprehensive survey of the Plasmodium life cycle by genomic, transcriptomic, and proteomic analyses. Science 2005, 307(5706):82–86.

25. Fougere A, Jackson AP, Bechtsi DP, Braks JA, Annoura T, Fonager J, Spaccapelo R, Ramesar J, Chevalley-Maurel S, Klop O et al Variant Exported Blood-Stage Proteins Encoded by Plasmodium Multigene Families Are Expressed in Liver Stages Where They Are Exported into the Parasitophorous Vacuole. PLoS Pathog 2016, 12(11):e1005917.

26. Otto TD, Bohme U, Jackson AP, Hunt M, Franke-Fayard B, Hoeijmakers WA, Religa AA, Robertson L, Sanders M, Ogun SA et al A comprehensive evaluation of rodent malaria parasite genomes and gene expression. BMC Biol 2014, 12:86.

27. Bafort JM The biology of rodent malaria with particular reference to Plasmodium vinckei vinckei Rodhain 1952. Ann Soc Belges Med Trop Parasitol Mycol 1971, 51(1):5–203.

28. Carter R, Walliker D New observations on the malaria parasites of rodents of the Central African Republic - Plasmodium vinckei petteri subsp. nov. and Plasmodium chabaudi Landau, 1965. Ann Trop Med Parasitol 1975, 69(2):187–196.

29. Carter R, Walliker D Malaria parasites of rodents of the Congo (Brazzaville): Plasmodium chabaudi adami subsp. nov. and Plasmodium vinckei lentum Landau, Michel, Adam and Boulard, 1970. Ann Parasitol Hum Comp 1976, 51(6):637–646.

30. Killick-Kendrick R Parasitic Protozoa of the blood of rodents. V. Plasmodium vinckei brucechwatti subsp. nov. A malaria parasite of the thicket rat, Thamnomys rutilans, in Nigeria. Ann Parasitol Hum Comp 1975, 50(3):251–264.

31. Landau I, Michel JC, Adam JP, Boulard Y The life cycle of Plasmodium vinckei lentum subsp. nov. in the laboratory; comments on the nomenclature of the murine malaria parasites. Ann Trop Med Parasitol 1970, 64(3):315–323.

32. Bafort J New isolations of murine malaria in Africa: Cameroon. In: 5th International Congress of Protozoology, New York City Abstracts of papers p343: 1977.

33. Bafort J Etude du cycle biologique du Plasmodium v. vinckei Rodhain 1952. Ann Soc Belge Méd Trop 1969, 49(6):533–628.

34. Bafort JM, Molyneux DH Liver infections with Plasmodium v. vinckei in hosts refractory to blood infection. Trans R Soc Trop Med Hyg 1971, 65(1):13.

35. Lainson FA Observations on the morphology and electrophoretic variation of enzymes of the rodent malaria parasites of Cameroon, Plasmodium yoelii, P. chabaudi and P. vinckei. Parasitology 1983, 86 (Pt 2):221–229.

36. Carter R Enzyme variation in Plasmodium berghei and Plasmodium vinckei. Parasitology 1973, 66(2):297–307.

37. LaCrue AN, Scheel M, Kennedy K, Kumar N, Kyle DE Effects of artesunate on parasite recrudescence and dormancy in the rodent malaria model Plasmodium vinckei. PLoS One 2011, 6(10):e26689.

38. Gautret P, Deharo E, Chabaud AG, Ginsburg H, Landau I Plasmodium vinckei vinckei, P. v. lentum and P. yoelii yoelii: chronobiology of the asexual cycle in the blood. Parasite 1994, 1(3):235–239.

39. Chandra R, Kumar S, Puri SK Plasmodium vinckei: infectivity of arteether-sensitive and arteether-resistant parasites in different strains of mice. Parasitol Res 2011, 109(4):1143–1149.

40. Killick-Kendrick R, Peters W Rodent malaria. London: Academic Press; 1978.

41. Kooij TW, Carlton JM, Bidwell SL, Hall N, Ramesar J, Janse CJ, Waters AP A Plasmodium whole-genome synteny map: indels and synteny breakpoints as foci for species-specific genes. PLoS Pathog 2005, 1(4):e44.

42. Carlton J, Angiuoli S, Suh B, Kooij T, Pertea M, Silva J, Ermolaeva M, Allen J, Selengut J, Koo H et al Genome sequence and comparative analysis of the model rodent malaria parasite Plasmodium yoelii yoelii. Nature 2002, 419(6906):512–519.

43. Liu SL, Sanderson KE Rearrangements in the genome of the bacterium Salmonella typhi. Proc Natl Acad Sci U S A 1995, 92(4):1018–1022.

44. Abkallo HM, Martinelli A, Inoue M, Ramaprasad A, Xangsayarath P, Gitaka J, Tang J, Yahata K, Zoungrana A, Mitaka H et al Rapid identification of genes controlling virulence and immunity in malaria parasites. PLoS Pathog 2017, 13(7):e1006447.

45. Akinosoglou KA, Bushell ES, Ukegbu CV, Schlegelmilch T, Cho JS, Redmond S, Sala K, Christophides GK, Vlachou D Characterization of Plasmodium developmental transcriptomes in Anopheles gambiae midgut reveals novel regulators of malaria transmission. Cell Microbiol 2015, 17(2):254–268.

46. Ukegbu CV, Giorgalli M, Tapanelli S, Rona LDP, Jaye A, Wyer C, Angrisano F, Blagborough AM, Christophides GK, Vlachou D PIMMS43 is required for malaria parasite immune evasion and sporogonic development in the mosquito vector. Proc Natl Acad Sci U S A 2020, 117(13):7363–7373.

47. Sultan AA, Thathy V, Frevert U, Robson KJ, Crisanti A, Nussenzweig V, Nussenzweig RS, Menard R: TRAP is necessary for gliding motility and infectivity of plasmodium sporozoites. Cell 1997, 90(3):511–522.

48. Sanders PR, Gilson PR, Cantin GT, Greenbaum DC, Nebl T, Carucci DJ, McConville MJ, Schofield L, Hodder AN, Yates JR, 3rd et al Distinct protein classes including novel merozoite surface antigens in Raft-like membranes of Plasmodium falciparum. J Biol Chem 2005, 280(48):40169–40176.

49. Taechalertpaisarn T, Crosnier C, Bartholdson SJ, Hodder AN, Thompson J, Bustamante LY, Wilson DW, Sanders PR, Wright GJ, Rayner JC et al Biochemical and functional analysis of two Plasmodium falciparum blood-stage 6-cys proteins: P12 and P41. PLoS One 2012, 7(7):e41937.

50. Ramaprasad A, Subudhi AK, Culleton R, Pain A A fast and cost-effective microsampling protocol incorporating reduced animal usage for time-series transcriptomics in rodent malaria parasites. Malar J 2019, 18(1):26.

51. Favaloro JM, Kemp DJ Sequence diversity of the erythrocyte membrane antigen 1 in various strains of Plasmodium chabaudi. Mol Biochem Parasitol 1994, 66(1):39–47.

52. Miller JL, Murray S, Vaughan AM, Harupa A, Sack B, Baldwin M, Crispe IN, Kappe SH Quantitative bioluminescent imaging of pre-erythrocytic malaria parasite infection using luciferase-expressing Plasmodium yoelii. PLoS One 2013, 8(4):e60820.

53. Franke-Fayard B, Trueman H, Ramesar J, Mendoza J, van der Keur M, van der Linden R, Sinden RE, Waters AP, Janse CJ A Plasmodium berghei reference line that constitutively expresses GFP at a high level throughout the complete life cycle. Mol Biochem Parasitol 2004, 137(1):23–33.

54. Boundenga L, Ngoubangoye B, Ntie S, Moukodoum ND, Renaud F, Rougeron V, Prugnolle F Rodent malaria in Gabon: Diversity and host range. Int J Parasitol Parasites Wildl 2019, 10:117–124.

55. Palmer GH, Brayton KA Gene conversion is a convergent strategy for pathogen antigenic variation. Trends Parasitol 2007, 23(9):408–413.

56. Subudhi AK, O’Donnell AJ, Ramaprasad A, Abkallo HM, Kaushik A, Ansari HR, Abdel-Haleem AM, Ben Rached F, Kaneko O, Culleton R et al Malaria parasites regulate intra-erythrocytic development duration via serpentine receptor 10 to coordinate with host rhythms. Nat Commun 2020, 11(1):2763.

57. Voza T, Gautret P, Renia L, Gantier JC, Lombard MN, Chabaud AG, Landau I Variation in murid Plasmodium desequestration and its modulation by stress and pentoxifylline. Parasitol Res 2002, 88(4):344–349.

58. Clark IA, Cowden WB, Butcher GA, Hunt NH Possible roles of tumor necrosis factor in the pathology of malaria. Am J Pathol 1987, 129(1):192–199.

59. Yoeli M, Hargreaves BJ Brain capillary blockage produced by a virulent strain of rodent malaria. Science 1974, 184(4136):572–573.

60. Tang J, Templeton TJ, Cao J, Culleton R The Consequences of Mixed-Species Malaria Parasite Co-Infections in Mice and Mosquitoes for Disease Severity, Parasite Fitness, and Transmission Success. Front Immunol 2019, 10:3072.

61. Abkallo HM, Tangena JA, Tang J, Kobayashi N, Inoue M, Zoungrana A, Colegrave N, Culleton R Within-host competition does not select for virulence in malaria parasites; studies with Plasmodium yoelii. PLoS Pathog 2015, 11(2):e1004628.

62. Martinelli A, Cheesman S, Hunt P, Culleton R, Raza A, Mackinnon M, Carter R A genetic approach to the de novo identification of targets of strain-specific immunity in malaria parasites. Proc Natl Acad Sci U S A 2005, 102(3):814–819.

63. Culleton R, Martinelli A, Hunt P, Carter R Linkage group selection: rapid gene discovery in malaria parasites. Genome Res 2005, 15(1):92–97.

64. Pattaradilokrat S, Culleton RL, Cheesman SJ, Carter R Gene encoding erythrocyte binding ligand linked to blood stage multiplication rate phenotype in Plasmodium yoelii yoelii. Proc Natl Acad Sci U S A 2009, 106(17):7161–7166.

65. Knowles G, Sanderson A, Walliker D Plasmodium yoelii: genetic analysis of crosses between two rodent malaria subspecies. Exp Parasitol 1981, 52(2):243–247.

66. Letunic I, Bork P Interactive Tree Of Life (iTOL) v4: recent updates and new developments. Nucleic Acids Res 2019, 47(W1):W256–W259.

67. Chin CS, Peluso P, Sedlazeck FJ, Nattestad M, Concepcion GT, Clum A, Dunn C, O’Malley R, Figueroa-Balderas R, Morales-Cruz A et al Phased diploid genome assembly with single-molecule real-time sequencing. Nat Methods 2016, 13(12):1050–1054.

68. Otto TD, Sanders M, Berriman M, Newbold C Iterative Correction of Reference Nucleotides (iCORN) using second generation sequencing technology. Bioinformatics 2010, 26(14):1704–1707.

69. Zerbino DR, Birney E Velvet: algorithms for de novo short read assembly using de Bruijn graphs. Genome Res 2008, 18(5):821–829.

70. Kurtz S, Phillippy A, Delcher AL, Smoot M, Shumway M, Antonescu C, Salzberg SL Versatile and open software for comparing large genomes. Genome Biol 2004, 5(2):R12.

71. Carver T, Harris SR, Berriman M, Parkhill J, McQuillan JA Artemis: an integrated platform for visualization and analysis of high-throughput sequence-based experimental data. Bioinformatics 2012, 28(4):464–469.

72. Carver TJ, Rutherford KM, Berriman M, Rajandream MA, Barrell BG, Parkhill J ACT: the Artemis Comparison Tool. Bioinformatics 2005, 21(16):3422–3423.

73. Robinson JT, Thorvaldsdottir H, Winckler W, Guttman M, Lander ES, Getz G, Mesirov JP Integrative genomics viewer. Nat Biotechnol 2011, 29(1):24–26.

74. Krzywinski M, Schein J, Birol I, Connors J, Gascoyne R, Horsman D, Jones SJ, Marra MA Circos: an information aesthetic for comparative genomics. Genome Res 2009, 19(9):1639–1645.

75. Stanke M, Waack S Gene prediction with a hidden Markov model and a new intron submodel. Bioinformatics 2003, 19 Suppl 2:ii215-225.

76. Kim D, Pertea G, Trapnell C, Pimentel H, Kelley R, Salzberg SL TopHat2: accurate alignment of transcriptomes in the presence of insertions, deletions and gene fusions. Genome Biol 2013, 14(4):R36.

77. Campbell MS, Holt C, Moore B, Yandell M Genome Annotation and Curation Using MAKER and MAKER-P. Curr Protoc Bioinformatics 2014, 48:4 11 11-14 11 39.

78. Lagesen K, Hallin P, Rodland EA, Staerfeldt HH, Rognes T, Ussery DW: RNAmmer: consistent and rapid annotation of ribosomal RNA genes. Nucleic Acids Res 2007, 35(9):3100–3108.

79. Jones P, Binns D, Chang HY, Fraser M, Li W, McAnulla C, McWilliam H, Maslen J, Mitchell A, Nuka G et al InterProScan 5: genome-scale protein function classification. Bioinformatics 2014, 30(9):1236–1240.

80. Krogh A, Larsson B, von Heijne G, Sonnhammer EL Predicting transmembrane protein topology with a hidden Markov model: application to complete genomes. J Mol Biol 2001, 305(3):567–580.

81. Petersen TN, Brunak S, von Heijne G, Nielsen H SignalP 4.0: discriminating signal peptides from transmembrane regions. Nat Methods 2011, 8(10):785–786.

82. Sargeant TJ, Marti M, Caler E, Carlton JM, Simpson K, Speed TP, Cowman AF Lineage-specific expansion of proteins exported to erythrocytes in malaria parasites. Genome Biol 2006, 7(2):R12.

83. Li H, Durbin R Fast and accurate short read alignment with Burrows-Wheeler transform. Bioinformatics 2009, 25(14):1754–1760.

84. Morgulis A, Gertz EM, Schaffer AA, Agarwala R A fast and symmetric DUST implementation to mask low-complexity DNA sequences. J Comput Biol 2006, 13(5):1028–1040.

85. McKenna A, Hanna M, Banks E, Sivachenko A, Cibulskis K, Kernytsky A, Garimella K, Altshuler D, Gabriel S, Daly M et al The Genome Analysis Toolkit: a MapReduce framework for analyzing next-generation DNA sequencing data. Genome Res 2010, 20(9):1297–1303.

86. Zhang Z, Li J, Zhao XQ, Wang J, Wong GK, Yu J KaKs_Calculator: calculating Ka and Ks through model selection and model averaging. Genomics Proteomics Bioinformatics 2006, 4(4):259–263.

87. Li L, Stoeckert CJ, Jr., Roos DS OrthoMCL: identification of ortholog groups for eukaryotic genomes. Genome Res 2003, 13(9):2178–2189.

88. Edgar RC MUSCLE: a multiple sequence alignment method with reduced time and space complexity. BMC Bioinformatics 2004, 5:113.

89. Capella-Gutierrez S, Silla-Martinez JM, Gabaldon T trimAl: a tool for automated alignment trimming in large-scale phylogenetic analyses. Bioinformatics 2009, 25(15):1972–1973.

90. Stamatakis A RAxML version 8: a tool for phylogenetic analysis and post-analysis of large phylogenies. Bioinformatics 2014, 30(9):1312–1313.

91. Larsson A AliView: a fast and lightweight alignment viewer and editor for large datasets. Bioinformatics 2014, 30(22):3276–3278.

