## Supplementary figures and images for "*Plasmodium vinckei* genomes provide insights into the pan-genome and evolution of rodent malaria parasites"

### AdditionalFile2

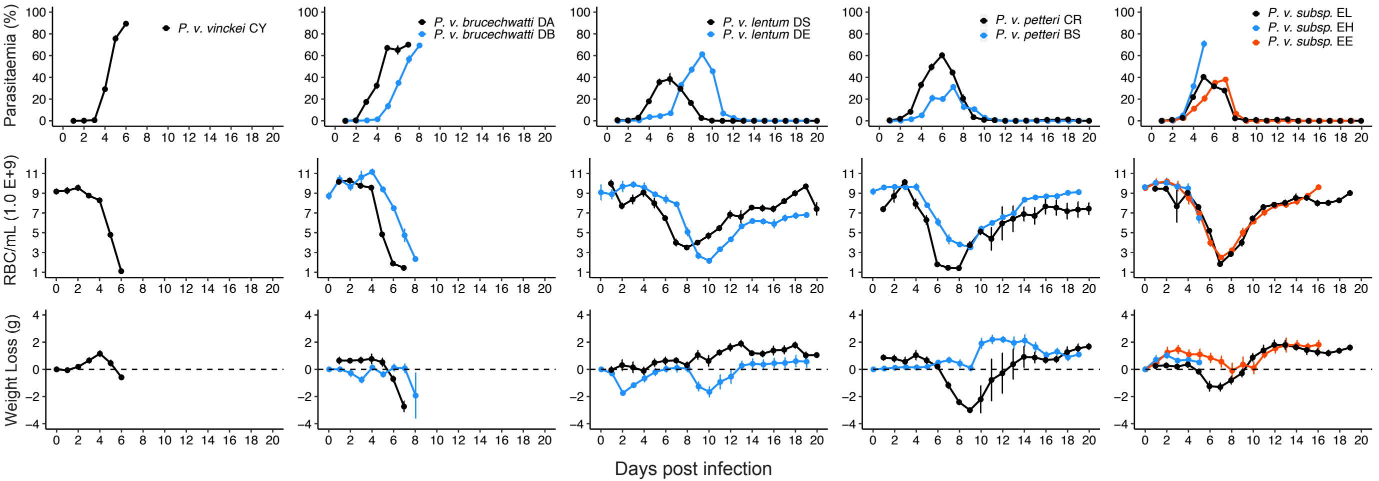

### AdditionalFile12

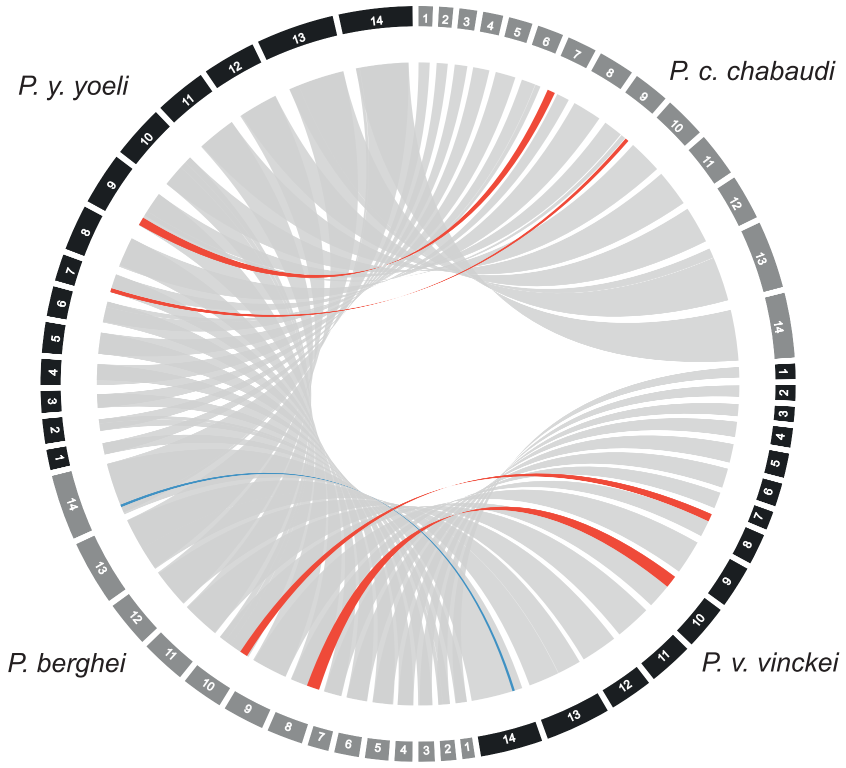

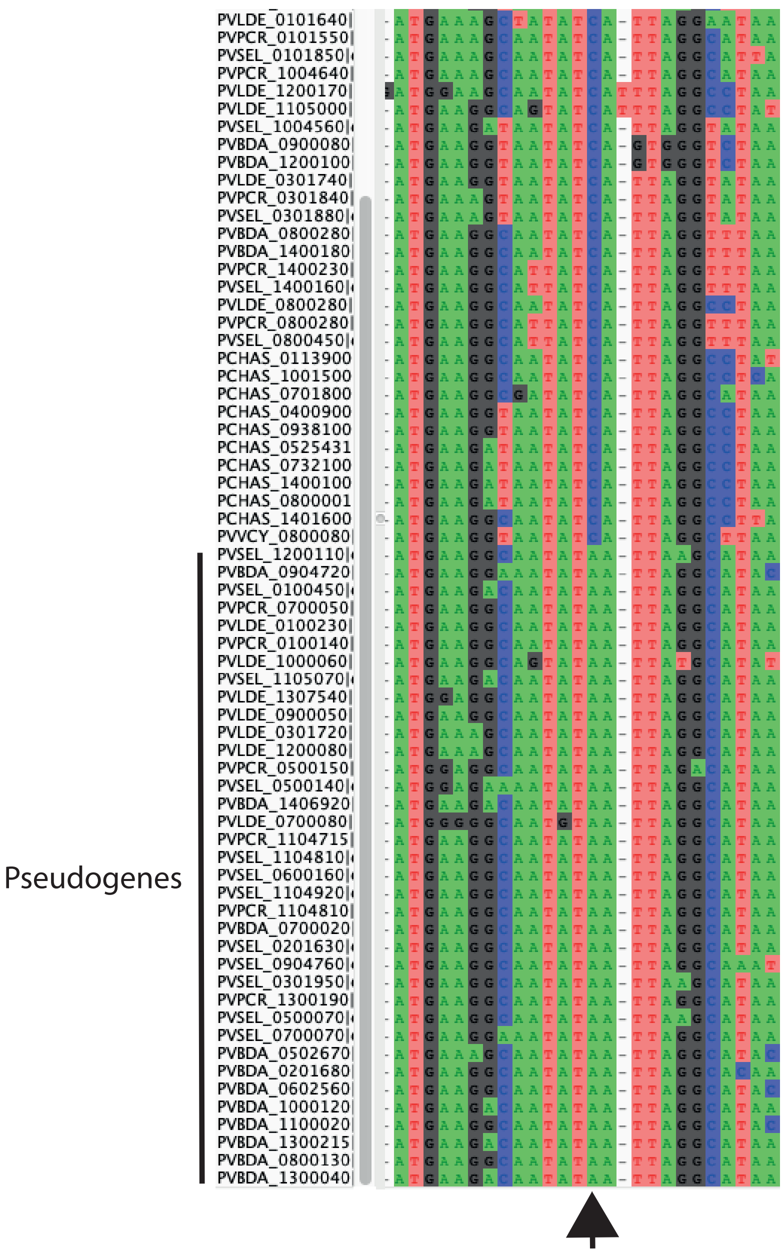
