## AdditionalFile9 for "*Plasmodium vinckei* genomes provide insights into the pan-genome and evolution of rodent malaria parasites"

### 1. erythrocyte membrane antigen 1 (*ema1*)

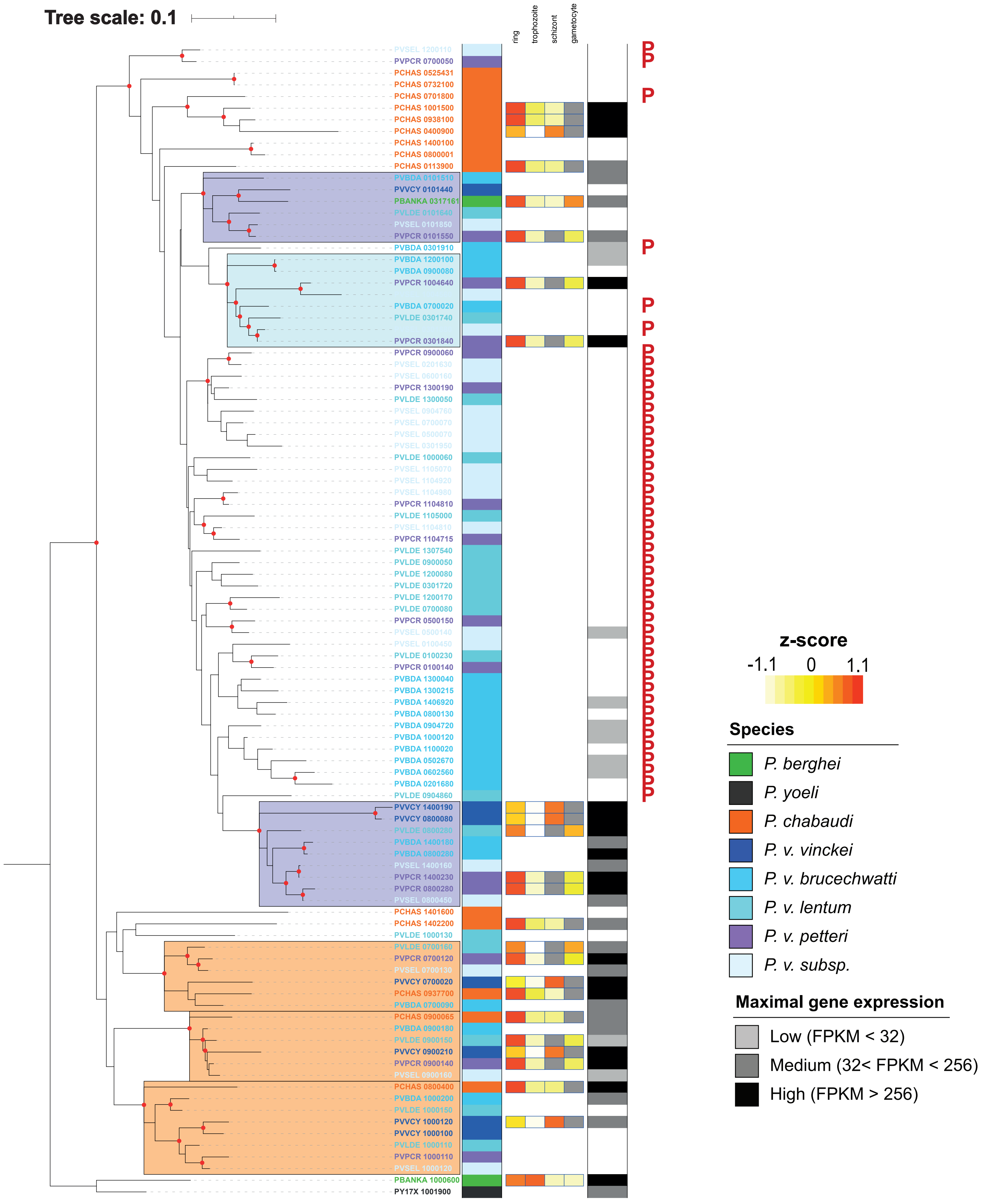

### 2. early transcribed membrane protein (*etramp*)

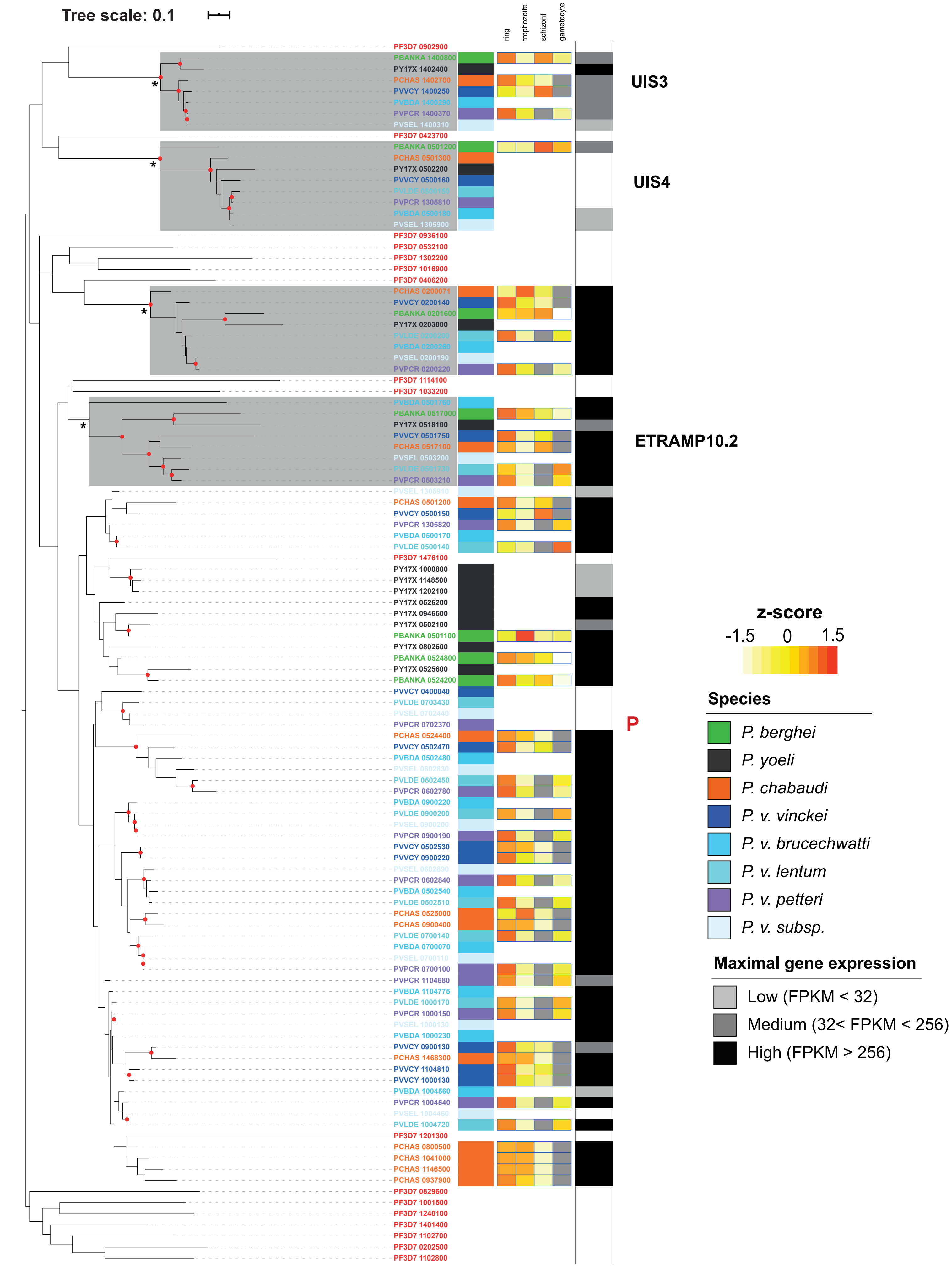

### 3. RMP-*fam-a*

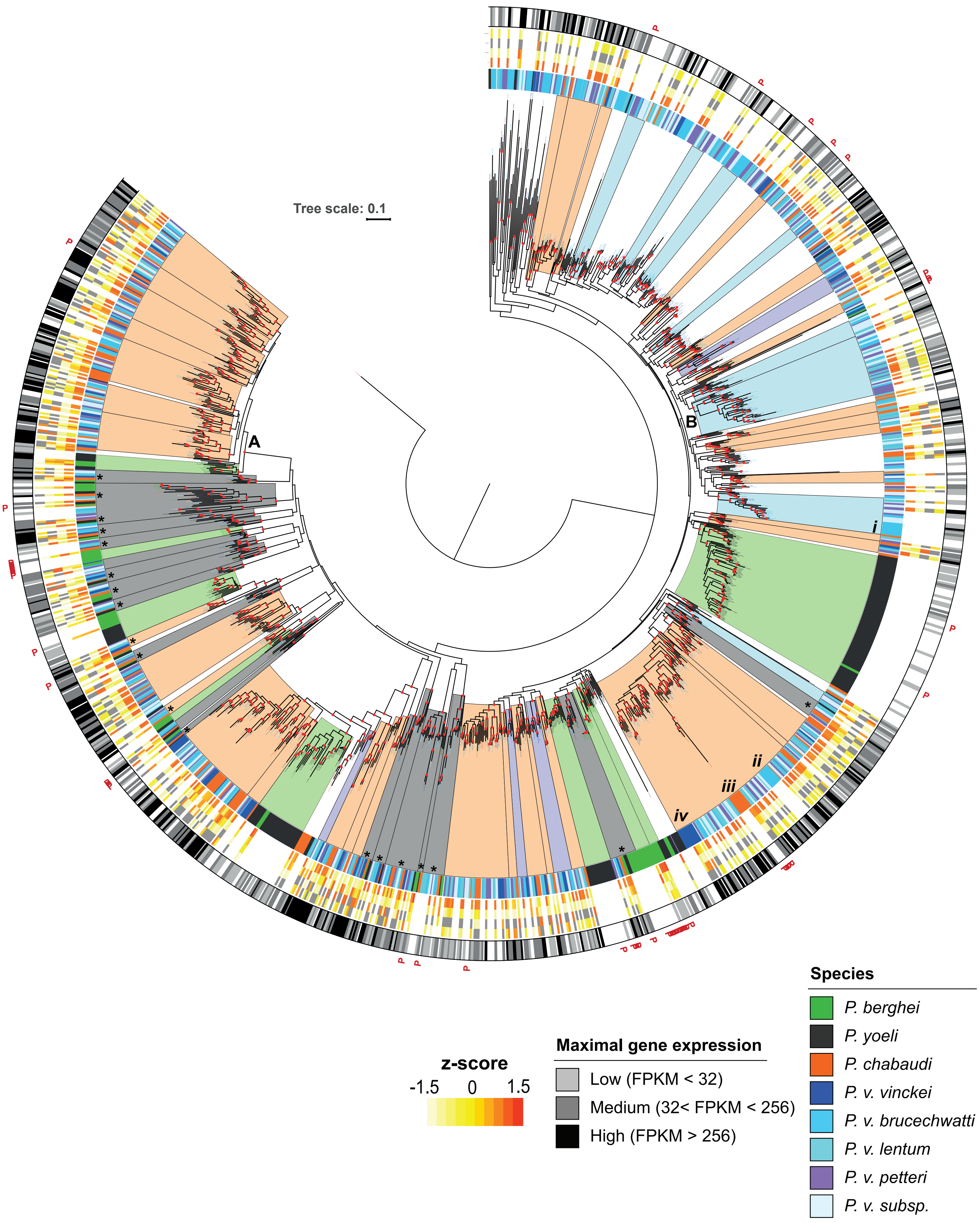
Tree scale: 0.1
A
B
i
ii
iii
iv
z-score
-1.5
0
1.5
Maximal gene expression
Low (FPKM < 32)
Medium (32< FPKM < 256)
High (FPKM > 256)
Species
P. berghei
P. yoeli
P. chabaudi
P. v. vinckei
P. v. brucechwatti
P. v. lentum
P. v. petteri
P. v. subsp.

### 4. RMP-*fam-b*

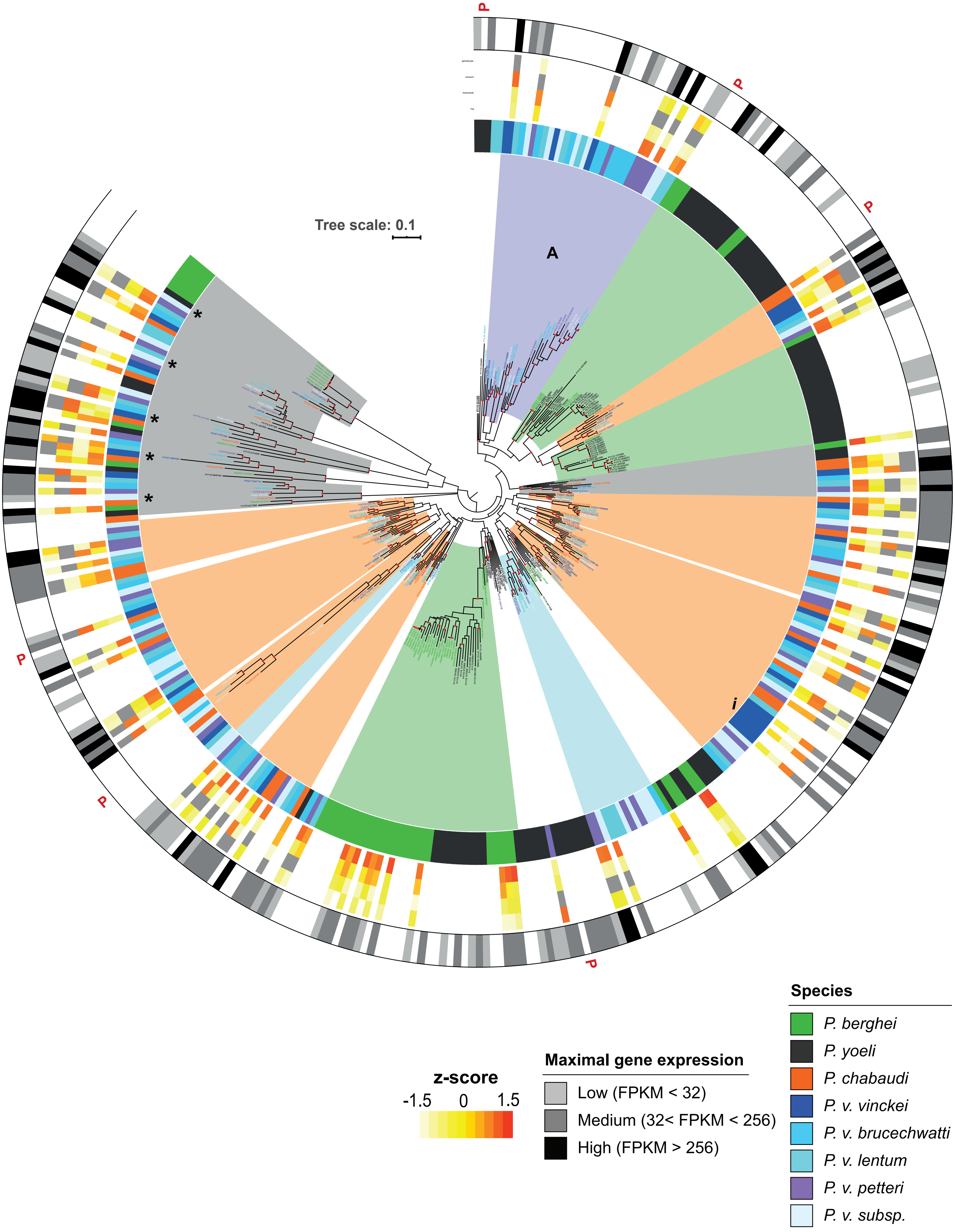

Tree scale: 0.1
A
*
*
*
*
*
i
P
P
P
P
P
P
z-score
-1.5
0
1.5
Maximal gene expression
Low (FPKM < 32)
Medium (32< FPKM < 256)
High (FPKM > 256)
Species
P. berghei
P. yoeli
P. chabaudi
P. v. vinckei
P. v. brucechwatti
P. v. lentum
P. v. petteri
P. v. subsp.

### 5. RMP-*fam-c*

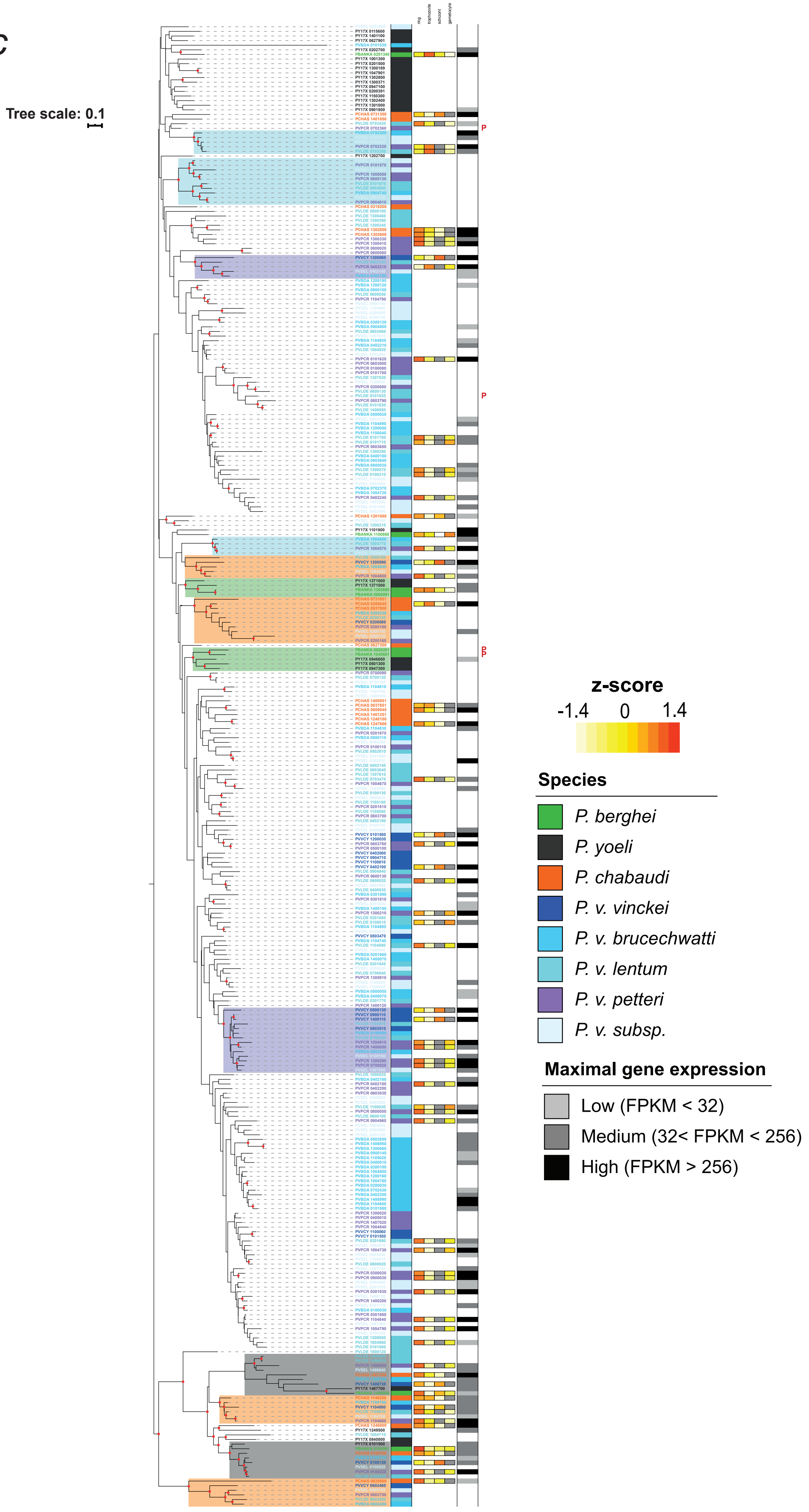

Tree scale: 0.1
ring
trophozoite
schizont
gametocyte
z-score
-1.4
0
1.4
Species
P. berghei
P. yoeli
P. chabaudi
P. v. vinckei
P. v. brucechwatti
P. v. lentum
P. v. petteri
P. v. subsp.
Maximal gene expression
Low (FPKM < 32)
Medium (32< FPKM < 256)
High (FPKM > 256)

### 6. RMP-*fam-d*

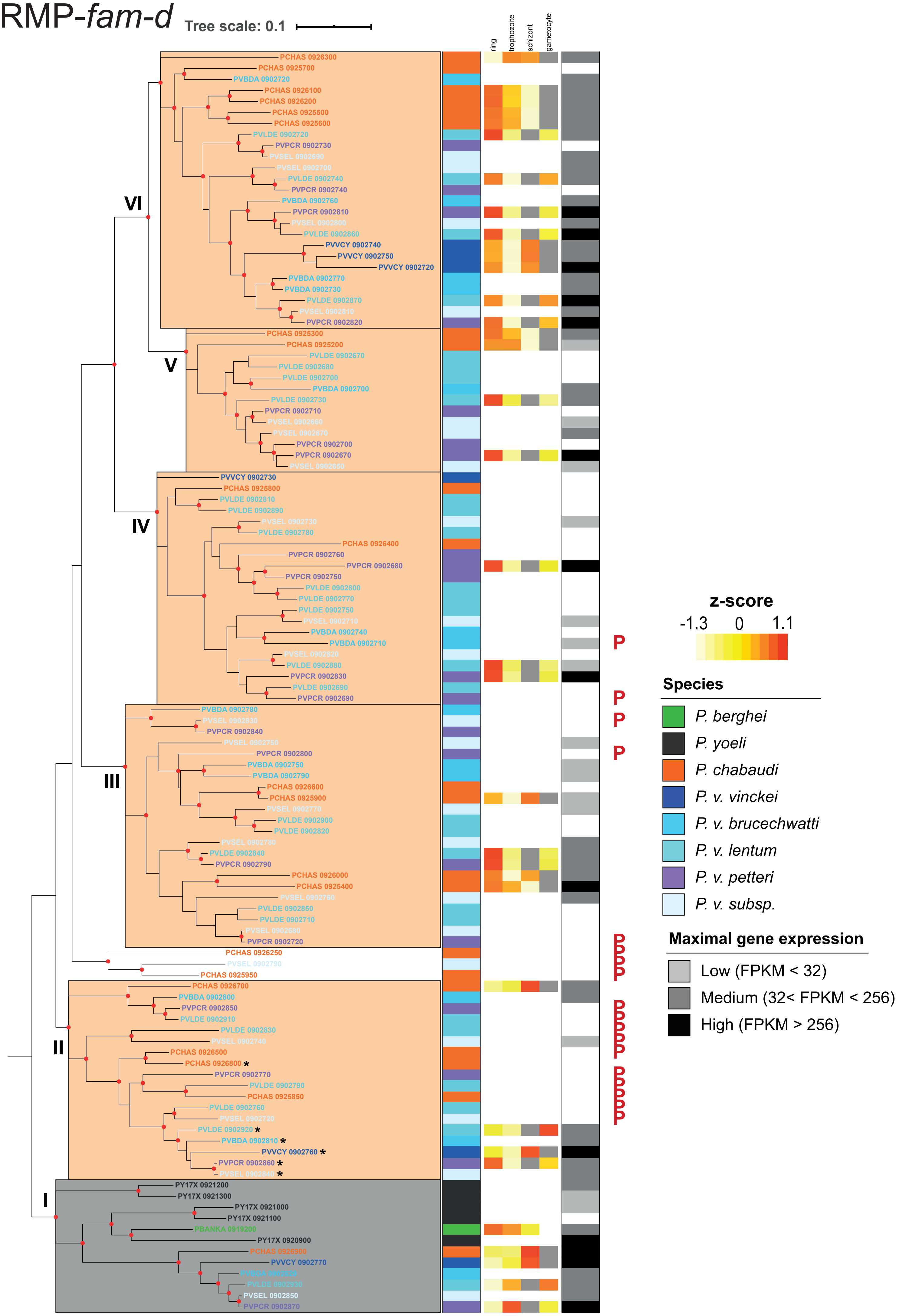

Tree scale: 0.1
ring
trophozoite
schizont
gametocyte
VI
V
IV
III
II
I
PCHAS 0926300
PCHAS 0925700
PVBDA 0902720
PCHAS 0926100
PCHAS 0926200
PCHAS 0925500
PCHAS 0925600
PVLDE 0902720
PVPCR 0902730
PVSEL 0902690
PVSEL 0902700
PVLDE 0902740
PVPCR 0902740
PVBDA 0902760
PVPCR 0902810
PVSEL 0902800
PVLDE 0902860
PVVCY 0902740
PVVCY 0902750
PVVCY 0902720
PVBDA 0902770
PVBDA 0902730
PVLDE 0902870
PVSEL 0902810
PVPCR 0902820
PCHAS 0925300
PCHAS 0925200
PVLDE 0902670
PVLDE 0902680
PVLDE 0902700
PVBDA 0902700
PVLDE 0902730
PVPCR 0902710
PVSEL 0902660
PVSEL 0902670
PVPCR 0902700
PVPCR 0902670
PVSEL 0902650
PVVCY 0902730
PCHAS 0925800
PVLDE 0902810
PVLDE 0902890
PVSEL 0902730
PVLDE 0902780
PCHAS 0926400
PVPCR 0902760
PVPCR 0902680
PVPCR 0902750
PVLDE 0902800
PVLDE 0902770
PVLDE 0902750
PVSEL 0902710
PVBDA 0902740
PVBDA 0902710
PVSEL 0902820
PVLDE 0902880
PVPCR 0902830
PVLDE 0902690
PVPCR 0902690
PVBDA 0902780
PVSEL 0902830
PVPCR 0902840
PVSEL 0902750
PVPCR 0902800
PVBDA 0902750
PVBDA 0902790
PCHAS 0926600
PCHAS 0925900
PVSEL 0902770
PVLDE 0902900
PVLDE 0902820
PVSEL 0902780
PVLDE 0902840
PVPCR 0902790
PCHAS 0926000
PCHAS 0925400
PVSEL 0902760
PVLDE 0902850
PVLDE 0902710
PVSEL 0902680
PVPCR 0902720
PCHAS 0926250
PVSEL 0902790
PCHAS 0925950
PCHAS 0926700
PVBDA 0902800
PVPCR 0902850
PVLDE 0902910
PVLDE 0902830
PVSEL 0902740
PCHAS 0926500
PCHAS 0926800 *
PVPCR 0902770
PVLDE 0902790
PCHAS 0925850
PVLDE 0902760
PVSEL 0902720
PVLDE 0902920 *
PVBDA 0902810 *
PVVCY 0902760 *
PVPCR 0902860 *
PVSEL 0902840 *
PY17X 0921200
PY17X 0921300
PY17X 0921000
PY17X 0921100
PBANKA 0919200
PY17X 0920900
PCHAS 0926900
PVVCY 0902770
PVLDE 0902930
PVSEL 0902850
PVPCR 0902870
P
P
P
P
PPPP
PPPPP
PPPPP
z-score
-1.3
0
1.1
Species
P. berghei
P. yoeli
P. chabaudi
P. v. vinckei
P. v. brucechwatti
P. v. lentum
P. v. petteri
P. v. subsp.
Maximal gene expression
Low (FPKM < 32)
Medium (32< FPKM < 256)
High (FPKM > 256)

### 7. haloacid dehalogenase-like hydrolases (*hdh*)

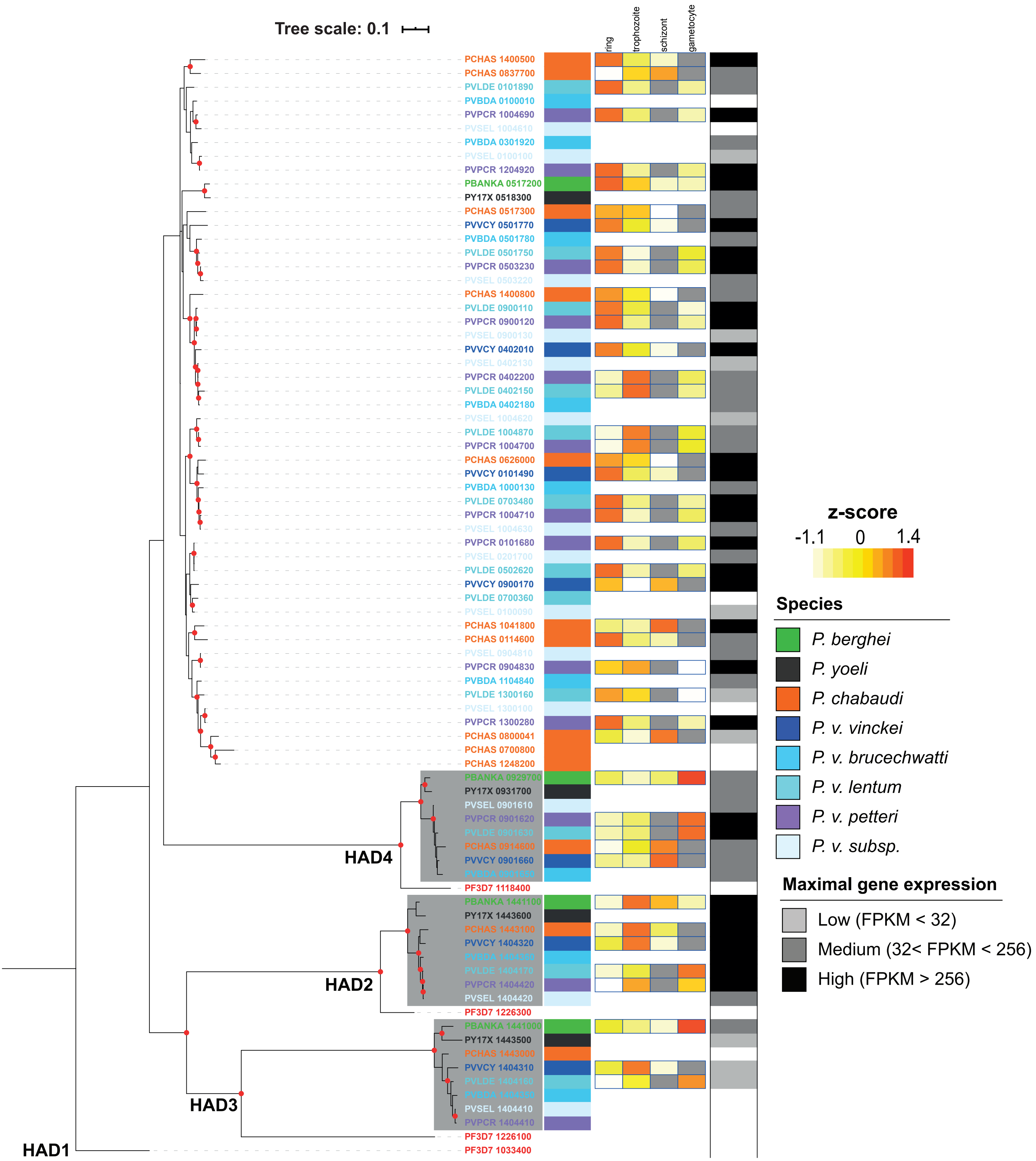

### 8. lysophospholipases (*lpl*)

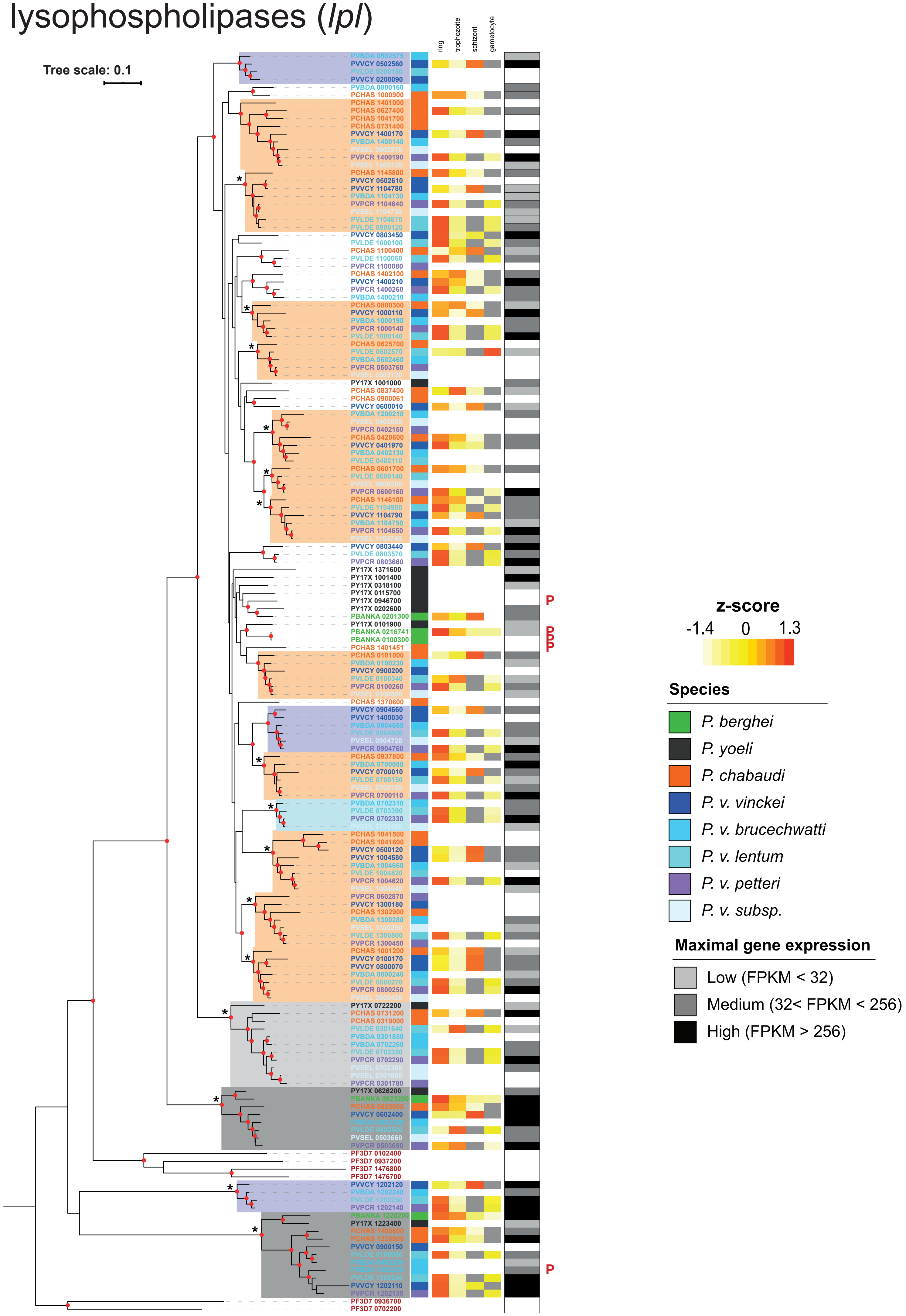

### 9. reticulocyte binding protein (*p235*)

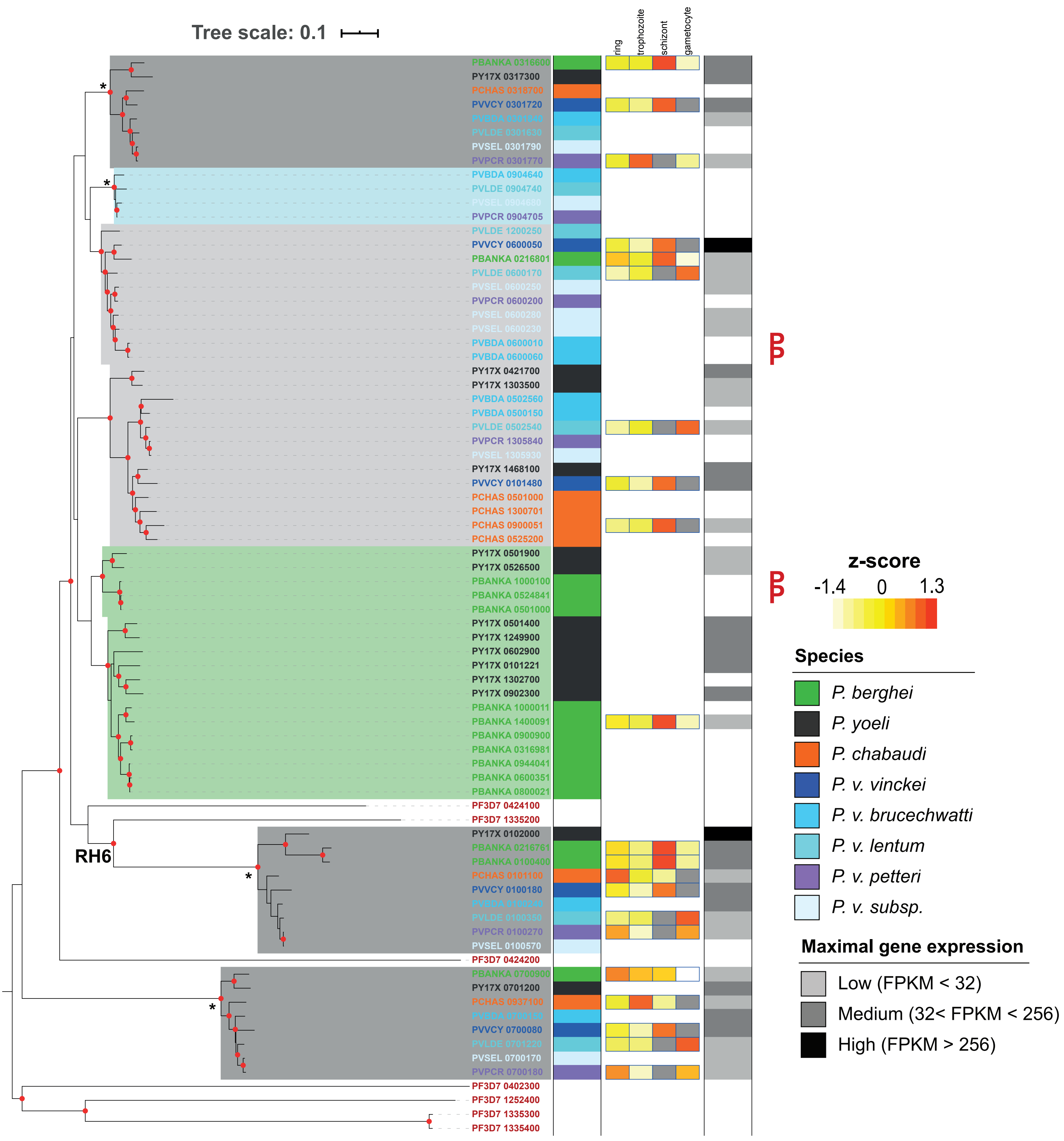

### 10. *Plasmodium* interspersed repeat proteins (*pir*)

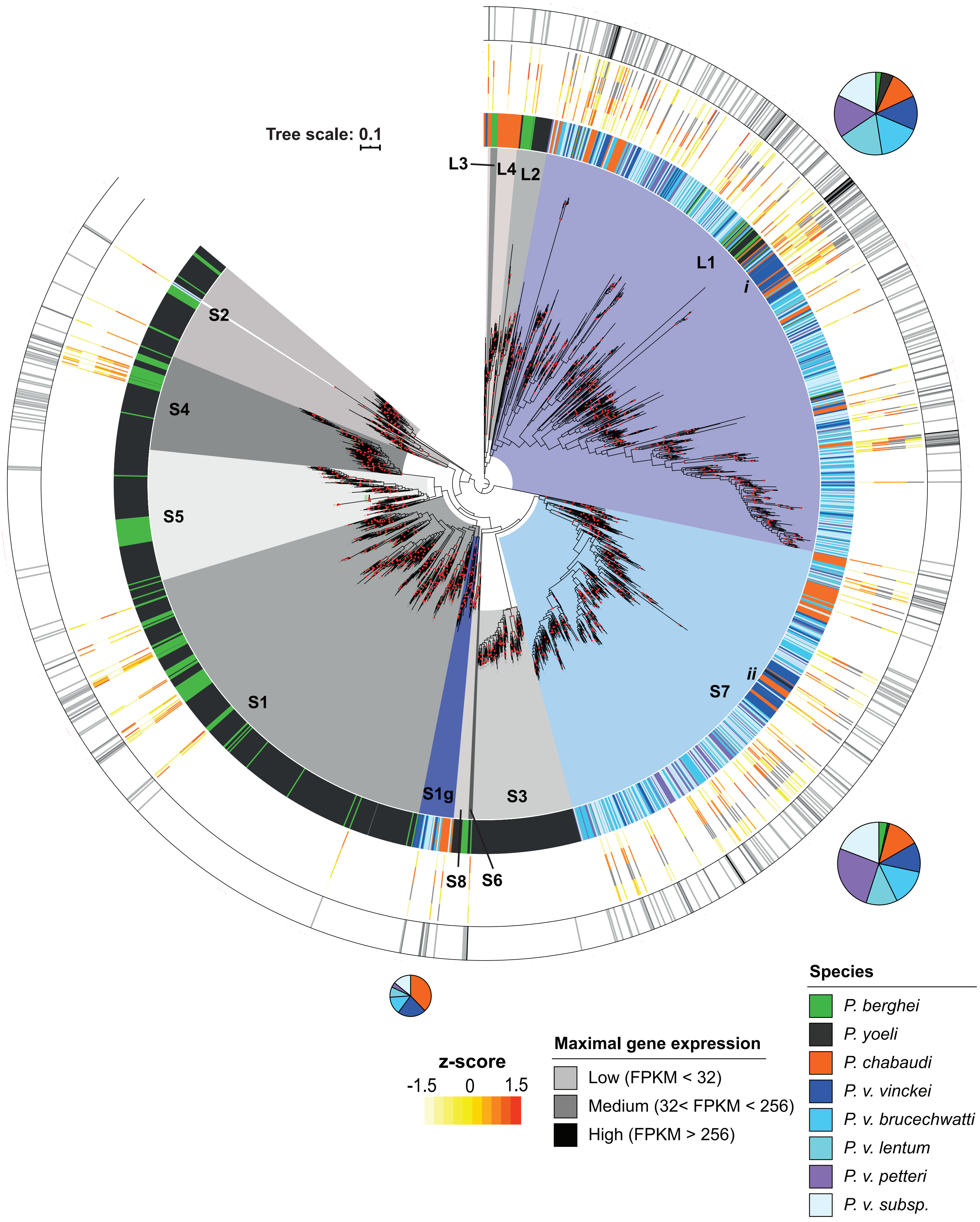
